# Sensitivity analysis of a mechanistic model of rumen fermentation and methane production by rumen microbiota in the presence of *Asparagopsis taxiformis*

**DOI:** 10.1101/2023.11.30.569127

**Authors:** Katarina Merk, Kathryn G. Link, Robert D. Guy, Matthias Hess

## Abstract

Ruminant animals rely on microbes for the conversion of complex plant material into host accessible metabolites. During this anaerobic conversion of plant biomass, termed enteric fermentation, methanogenic archaea convert hydrogen into the potent greenhouse gas methane (CH_4_). The search for methane mitigation strategies to combat climate change has identified the red seaweed *Asparagopsis taxiformis* as a promising feed additive that, when added to a regular cattle diet, reduced enteric CH_4_ by over 80%. A more complete understanding of microbial interactions during enteric fermentation is needed for ongoing improvement to mitigation methods. Mathematical models that permit *in silico* simulation of enteric fermentation allow for the identification of key parameters that drive rumen methane production. Here we built upon an existing rumen fermentation model and calibrated it using a preliminary classification of functional microbial groups and gas emission data from a previously published *in vitro* rumen fermentation experiment, but many microbes remained functionally unclassified. The model was then used to conduct an *in silico* experiment to explore how the partition of functionally unclassified microbes into functional groups affects methane output. These *in silico* experiments identified that model methane production is more sensitive to microbial variation in the presence of *A. taxiformis* versus without. The use of local and global sensitivity analysis approaches revealed other rumen parameters to also be drivers of enteric methane production. In the presence of *A. taxiformis*, parameters modulating methane production include bromoform concentration, methanogen abundance, total microbial concentration, a parameter effecting the inhibition of methanogen growth rate by the action of bromoform, and the maximum specific utilization rate of hydrogen. Without *A. taxiformis*, feed composition parameters, the hydrolysis rate constant of cell wall carbohydrates, and a parameter affecting the yield factors during sugar utilization were found to be most significant. For possible methane reduction without *A. taxiformis*, we propose an adjustment in feed composition parameters that reduces predicted methane by 25.6%.

## Introduction

Ruminant animals (cattle, goats, sheep, etc.) are vital sources of both food and income but can produce between 250 and 500 liters of methane daily and thus are substantial contributors to global warming (1). This ruminant-derived methane (CH_4_) has a global warming potential that is 28-fold higher than carbon dioxide on a 100-year scale (2). Additionally, the eructation of CH_4_ has been estimated to account for 8-12% direct digested energy loss for the ruminant and results in decreased economic gains for producers (1), further supporting the need for strategies to reduce enteric CH_4_.

Feed additives are one potential solution, with the red seaweed *Asparagopsis taxiformis* emerging as a promising candidate for significant methane reduction in ruminants (3). The methane reduction powers of *A. taxiformis* have been attributed to the bromoform content of this seaweed and its ability to inhibit the key enzyme of archaeal methanogenesis, but the complete mode-of-action of this bromoform-rich red seaweed is still not fully understood.

The rumen ecosystem is a complex network of microbes that work synergistically to break down and ferment feed in the digestive system. Understanding, and subsequently manipulating, this microbiome has the potential to improve feed efficiency and reduce methane emissions. Even though there is a strong connection between rumen microbiota and methane emissions, success of microbial manipulation to address global agricultural challenges has been limited (4). There have been significant advancements in omics techniques that have expanded our understanding of the rumen microbiota consortia and function, but applying this knowledge for microbial manipulation remains a challenge.

The diversity of the rumen microbiome and complexity of interactions in rumen fermentation make it difficult to understand the behavior of this gut ecosystem without thorough experimentation methods and companion quantitative methods. To address this need for quantitative methods, a growing number of mathematical models have been developed to improve our understanding of rumen fermentation (5–16). The main structures of the dynamic models are built upon Baldwin et al. (5) and Dijkstra et al. (6). Such models can be powerful tools as they allow tracking of individual state variables during rumen fermentation simulations, something less feasible with *in vitro* or *in vivo* methods. In addition, these models can be very helpful for interpreting experimental data, elucidating biochemical mechanisms, and guiding experimental design when properly calibrated, validated, and evaluated. However, current rumen fermentation models fail to incorporate microbial genomic data, which is necessary to improve applicability of the models and for enhanced understanding of rumen fermentation (14). For a current review on how microbial genome data can be incorporated in these mathematical models, see Munoz-Tamayo et al. (17). Models developed by Muñoz-Tamayo and van Lingen include the dynamics of methanogens (10–13), but they do not incorporate microbial genomic data in their analysis and thus could be further improved with the addition of this data type.

Studying the effects of methane inhibitors *in silico* can be helpful to improve understanding of their effects on rumen fermentation, and in particular the rumen microbiome. In addition to the dynamics of methanogens, Muñoz-Tamayo et al. (12) and van Lingen et al. (11) models include the effects of methane inhibitors. Of particular interest to the work here, Muñoz-Tamayo et al. (12) incorporates the effects of feed additive *A. taxiformis*. The effect of *A. taxiformis* is described in the model through the inclusion of direct methanogenesis inhibition by *A. taxiformis* component bromoform, indirectly through the hydrogen control of sugar utilization and volatile fatty acid (VFA) production by re-routed hydrogen. While these are all valid approaches to understanding the effect of *A. taxiformis* on methane production, a complete picture of rumen fermentation and methane output necessitates the incorporation of microbial data. Nevertheless, the Muñoz-Tamayo et al. (12) and van Lingen et al. (11) models serve as excellent starting points for the development of improved quantitative methods in the rumen fermentation field.

As was done with the Muñoz-Tamayo et al. (12) and van Lingen et al. (11) models, the values of rate constants utilized in these mathematical models are often measured in isolation, i.e., considering only one reaction and the involved reactants, and these experiments are conducted under varying conditions e.g., temperature. Thus, there is uncertainty in parameter values and how changes in parameter values affect the system. Understanding how model output changes when key parameters and variables vary is crucial in assessing the robustness of model predictions, and subsequently in applying the model findings to *in vitro* and *in vivo* systems.

Conducting sensitivity analysis (SA) on rumen fermentation models can help identify drivers of methane production. SA is the study of how the uncertainty of the model input attributes to uncertainty in the model output (18,19). SA helps determine parameters that dominate high-dimensional system behavior, providing potential targets for further downstream analysis. There are two main types of sensitivity analysis: local and global. Local sensitivity analysis (LSA) analyzes the impact of one parameter on the model output, while all other parameters remain constant. Global sensitivity analysis (GSA) examines the effect of multiple, simultaneous variations in model parameters on model output. LSA is straightforward in its application and interpretation, while GSA can be more involved with multiple approaches. SA has been conducted on various models of rumen fermentation (8–11,20–22), but has not been conducted in a model that incorporates the effects of *A. taxiformis* or that incorporates microbial data.

Here we describe our adaption and further calibration of the Muñoz-Tamayo et al. (12) model and the ensuing comparison to Roque et al. (3) continuous *in vitro* RUSITEC experimental data. The addition of rumen microbial community information from this experiment allowed us to explore how functional classifications of microbes affect the fermentation process, with particular focus on methane production. Though making some concessions for biological processes, to create a robust mathematical model we operate under the premise that it is feasible to attribute specific functionality to microbes, and that each microbe is exclusively involved in a single macroscopic reaction. These reductive microbes are then grouped into microbial functional groups (sugar utilizers, amino acid utilizers, and methanogens), which is comprised of various microbial species. We performed virtual microbiome experiments to study the importance of functional classifications as they pertain to methane output in the rumen ecosystem. Our *in silico* experiments showed that simulated methane production is significantly more sensitive to microbial variation with bromoform in the system than with no bromoform added. The optimal distribution of functionally unclassified microbes was found that best described methane production in Roque et al. (3).

We use local and global sensitivity analysis methods to identify key rumen parameters driving methane production. In the presence of *A. taxiformis*, parameters modulating methane production include bromoform concentration, methanogen abundance, total microbial concentration, a parameter affecting the inhibition of methanogen growth rate by the action of bromoform, and the maximum specific utilization rate of hydrogen. Without *A. taxiformis*, feed composition parameters, the hydrolysis rate constant of cell wall carbohydrates, and a parameter affecting the yield factors during sugar utilization were found to be most significant. Using the parameters identified in the sensitivity analysis, we propose feeding strategies for methane reduction in the absence of *A. taxiformis*. Here we highlight that the use of mathematical models in complex biological systems, such as the rumen, may reduce costly exploratory *in vivo* experimentation and provide scientific roadmaps to achieving desired biological outcomes.

## 1 Materials and Methods

### 1.1 Experimental Data

Experimental data from Roque et al. (3), an *in vitro* study of the impact of *A. taxiformis* on fermentation and methane production, was used for model calibration. The study showed a 95% reduction in methane production when *A. taxiformis* was added to the fermentation vessels at a 5% inclusion rate. The semi-continuous rumen simulation technique system (RUSITEC) was used. Rumen fluid and gas samples were collected 4, 12, and 24 h after vessel feeding every day. Carbon dioxide (CO_2_) and methane (CH_4_) concentrations were measured using an SRI Gas Chromatograph (8610C, SRI, Torrance, CA), gas volume was measured using miligas flow meter (Ritter, Germany), microbial data was assessed using high-throughput 16S ribosomal RNA gene amplicon sequencing and analyzed using mothur (23), and VFA profiles were determined using Gas Chromatography-Flame Ionization detection on a Thermo Trace GC Ultra machine (Thermo Electron Corporation, Rodano Milan, Italy). For model calibration, we considered the average data from vessels 1-3 for control and 4-6 for treatment.

In Roque et al. (3) rumen fluid and solids were collected from two independent cows, mixed, then divided into two vessels, one with *A. taxiformis* and one without. Super basic ration feed was added to both vessels, which were then allowed to adjust for 24 hours to let the systems reach a more consistent state before beginning the experiment. After this acclimation period, rumen fluid with and without *A. taxiformis* were each divided into 3 vessels and each vessel received one concentrate bag from their equilibration vessel in addition to a new concentrate bag. In the experiment, each vessel was opened every 24 hours, a feedbag and the gas bag were exchanged, and the vessels were flushed with nitrogen to maintain anaerobic conditions. To account for these discontinuities due to sampling time, integration was stopped at the time points of discontinuity (24 hour, 48 hour, 72 hour), initial conditions were set, and vessel attachment and integration was restarted. Initial conditions of the gaseous phase and feed variables were reset (see Supplement A for more detail) and the initial conditions of the remaining variables were set to the last value of the previous integration of their variable. Each concentrate bag remained in the vessel for 48 hours before exchange. The overlap allows the fiber adherent microbes to transfer from one feed substrate to the other, as is typical *in vivo*. To further mimic the *in vivo* system and maintain system pH, artificial saliva buffer was delivered to each vessel via peristaltic pump at 0.39mL/min throughout days 1-4.

### 1.2 Mathematical Modeling

We adapted and calibrated the Muñoz-Tamayo et al. (12) mathematical model of *in vitro* rumen fermentation with the Roque et al. (3) gas and microbial phylogenetic data from the *in vitro* semi-continuous RUSITEC system. We then conducted sensitivity analysis and identified rumen parameters that might be key drivers in enteric methane production. Details of the model assumptions, modifications, and calibration are described below.

The model assumes the rumen ecosystem is made up of four major component groups: particulate components, soluble components, gaseous phase components, and functional microbial groups (see Figure 1). The particulate components include fiber carbohydrates, non-fiber carbohydrates, and proteins. The soluble components include glucose, amino acids, ammonia (NH_3_), acetate, butyrate, propionate, CO_2_, H_2_, and CH_4_. The gaseous phase components include CO_2_, H_2_, and CH_4_. The functional microbial groups include sugar utilizers, amino acid utilizers, and methanogens. The breakdown of carbohydrates releases sugars and the breakdown of proteins releases amino acids, which either form products (soluble components) or are utilized and form microbial biomass. To capture the main function of the microbes in the fermentation process, microbes are classified as sugar utilizers, amino utilizers, and methanogens. The functional microbial group of sugar utilizers utilizes glucose and NH_3_, the amino acid utilizers metabolize amino acids, and the methanogens convert H_2_ to produce CH_4_. Mass transfer phenomena of CO_2_, CH_4_, H_2_ and occur between liquid and gas phases. *A. taxiformis* contains bromoform, which affects some of these interactions through direct inhibition of methane production and indirectly through hydrogen control.

**Figure 1:**
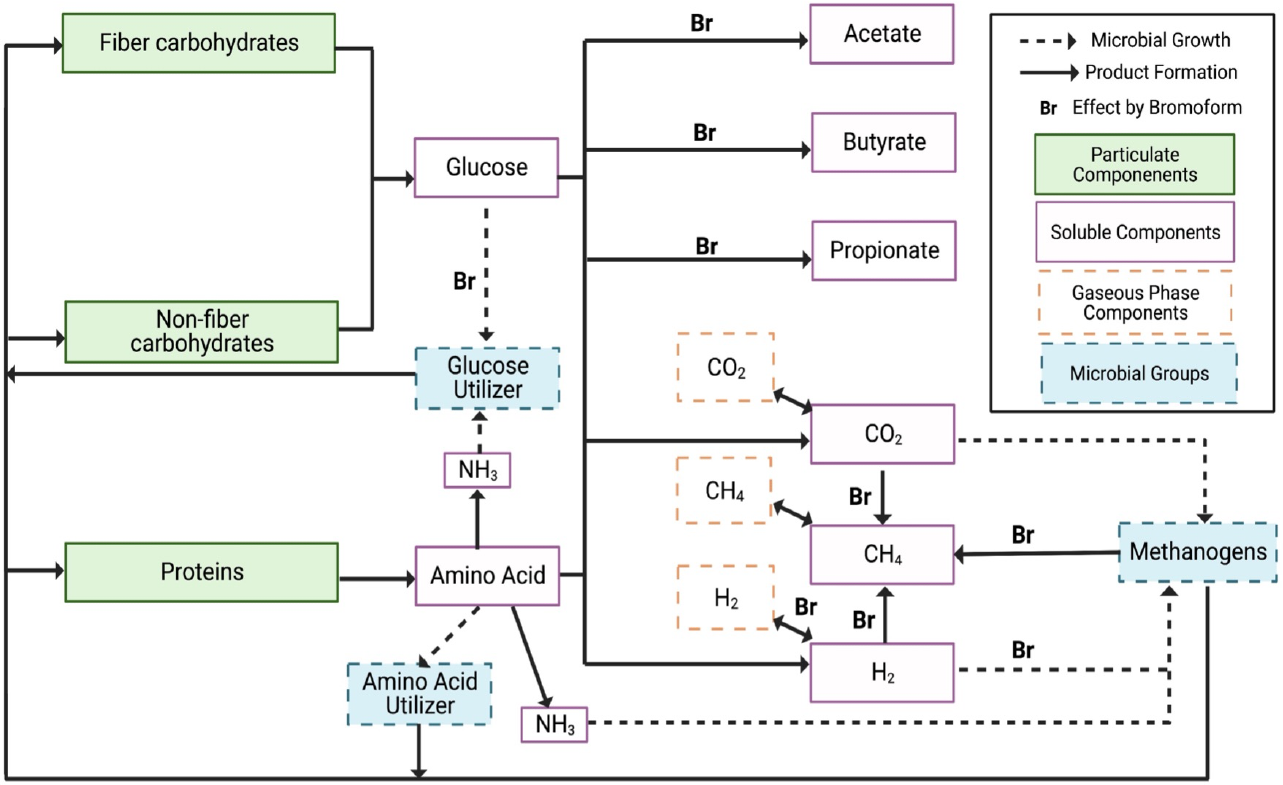
Rumen fermentation state diagram. Represents interaction in the rumen fermentation model (adapted from Muñoz-Tamayo et al (12)). The rumen ecosystem is made up of four major component groups: particulate components (solid green rectangles, green interiors), soluble components (solid purple rectangles, white interiors), gaseous phase components (dashed orange rectangles, white interiors), and functional microbial groups (dashed blue rectangles, blue interiors). Solid arrows indicate product formation and dashed arrows indicate microbial growth. Bromoform (**Br**) affects some of these interactions through direct inhibition of methane production and indirectly through hydrogen control.

The mathematical model contains 20 temporarily varying ordinary differential equations and over 50 parameters including reaction rates and initial conditions. To align our *in silico* experiments with Roque et al. (3), we extended the simulation time to 96 hours compared to 24 hours reported in Muñoz-Tamayo et al. 2021. The following model adaptions were made to the Muñoz-Tamayo et al.(12)model:

1. Microbial relative abundance data from Roque et al. (3) was used for determining initial conditions of microbial concentrations in the model. Muñoz-Tamayo et al. (13) and Muñoz-Tamayo et al (12) did not have microbial relative abundance data and used approximate values. With the use of the microbial relative abundance data, the importance of the rumen microbiome and its functional classifications was studied. Further detailed in section 2.3.2.
2. Separate liquid-gas transfer constants, *kLa*_2_ and *kLa*_3_were added for CO_2_ and H_2_ respectively, as different molecules evaporate at different rates and the addition of separate terms improved model fit.
3. Liquid dilution terms were added due to the continuous addition of buffer to the RUSITEC system in Roque et al. (3). Specifically, the total liquid volume was tracked as a function of time. The Muñoz-Tamayo et al. 2021 model assumed constant liquid volume given the experimental setup used for model calibration.
4. Volume in gas phase (*V*_*g*_) was calculated using linear best fit of volume data versus time. The Muñoz-Tamayo et al. 2021 model assumed constant gaseous volume. In Roque et al. (3) gaseous volume was zero at every feeding (due to exchange of feedbags) and increased for 24 hours after every feeding.

Each simulation with the mathematical model requires setting numerous input values. We specify the initial concentrations of all model species, the rate constants for all reactions, biochemical and physiochemical parameters. The outputs of the model are the concentrations of polymer, soluble, inorganic, gaseous, and microbial species at each instant of time including bromoform concertation. A complete listing of the model’s differential equations of the base parameter values used in the simulation can be found in S1 Appendix. A list of abbreviations used is in Table 1. Parameters, initial conditions, and their values are in Table 2.

**Table 1:** Model variable and parameter definitions. (Asterisk indicate parameters varied in sensitivity analysis. Asterisk next to state variable indicates that its initial condition was varied)

| Parameter | Definition | Units |
| --- | --- | --- |
| **Polymer components** |  |  |
| $Z_{ndf} *$ | Neutral detergent fiber concentration | g/L |
| $Z_{nsc} *$ | Non-structural carbohydrate concentration | g/L |
| $Z_{pro} *$ | Proteins concentration | g/L |
| **Soluble components** |  |  |
| $S_{su}$ | Sugars concentration | mol/L |
| $S_{aa}$ | Average amino acid concentration | mol/L |
| $S_{ac}$ | Total acetate concentration | mol/L |
| $S_{bu}$ | Total butyrate concentration | mol/L |
| $S_{pr}$ | Total propionate concentration | mol/L |
| $S_{IC} *$ | Inorganic carbon concentration | mol/L |
| $S_{IN}$ | Inorganic nitrogen concentration | mol/L |
| $S_{H_2}$ | Hydrogen concentration in liquid phase | mol/L |
| $S_{CH_4}$ | Methane concentration in liquid phase | mol/L |
| $S_{br}$ | Bromoform concentration | g/L |
| $V$ | Volume in liquid phase | L |
| **Gas components** |  |  |
| $n_{g,H_2}$ | Moles of hydrogen in gas phase | mol |
| $n_{g,CO_2}$ | Moles of carbon dioxide in gas phase | mol |
| $n_{g,CH_4}$ | Moles of methane in gas phase | mol |
| **Microbial functional groups** |  |  |
| $x_{su}$ | Concentration of sugar utilizing microbes | mol/L |
| $x_{aa}$ | Concentration of amino acid utilizing microbes | mol/L |
| $x_m$ | Concentration of methanogens | mol/L |
| $p_{su,0} *$ | Initial relative abundance of sugar utilizing microbes | |
| $p_{aa,0} *$ | Initial relative abundance of amino acid utilizing microbes | |
| $p_{m,0} *$ | Initial relative abundance of methanogens | |
| $TMC *$ | Total Microbial Concentration | mol/L |
| **Rates** |  |  |
| $\rho_i$ | Kinetic rate of microbial process j | mol (or g) j/(L h) |
| $\rho_{x_j}$ | Death cell rate of microbes j | mol j/(L h) |
| $\rho_{T_j}$ | Liquid-gas transfer rate of component j | mol j/(L h) |
| $\rho_b$ | Rate of saliva buffer | ml/min |
| $k_{br} *$ | Kinetic rate constant of bromoform utilization | 1/h |
| **Biochemical parameters** |  |  |
| $\lambda_k$ | Molar fraction of the sugars utilized via reaction k | mol/mol |
| $\sigma_{i,x}$ | Fraction of i of the biomass | g/g |
| $\sigma_{i,aa}$ | Stoichiometric coefficient of i production from amino acid fermentation | |
| $k_d$ | Death cell rate constant | $h^{-1}$ |
| $k_{hyd,ndf}$ * | Hydrolysis rate constant of cell wall carbohydrates | $h^{-1}$ |
| $k_{hyd,nsc}$ * | Hydrolysis rate constant of non-structural carbohydrates | $h^{-1}$ |
| $k_{hyd,pro}$ * | Hydrolysis constant of proteins | $h^{-1}$ |
| $k_{m,aa}$ * | Maximum specific utilization rate of amino acids | mol/(mol h) |
| $k_{m,H_2}$ * | Maximum specific utilization rate of hydrogen | mol/(mol h) |
| $k_{m,su}$ * | Maximum specific utilization rate of sugars | mol/(mol h) |
| $K_{s,aa}$ | Monod constant associated with the utilization of amino acid | mol/L |
| $K_{s,H_2}$ | Monod constant associated with the utilization of hydrogen | mol/L |
| $K_{s,su}$ | Monod constant associated with the utilization of sugars | mol/L |
| $K_{s,IN}$ * | Nitrogen limitation constant | mol/L |
| $Y_j$ * | Microbial biomass yield factor of j | mol/mol |
| $Y_{i,j}$ | Yield factor of the compound i during utilization of substrate j | mol/mol |
| **Physiochemical parameters** |  |  |
| $K_{a,CO_2}$ | Equilibrium constant of bicarbonate | |
| $kLa_i$ * | Liquid gas transfer constant for component i | $h^{-1}$ |
| $K_{H,CO_2}$ | Henry's law coefficient of carbon dioxide | M/bar |
| $K_{H,CH_4}$ | Henry's law coefficient of methane | M/bar |
| $K_{H,H_2}$ | Henry's law coefficient of hydrogen | M/bar |
| $p_{H_2}$ | Partial pressure of hydrogen | bars |
| $T$ | Temperature | K |
| $t_{full}$ | Time when vessels reach full capacity | h |
| $w_j$ | Molecular weight of component j | g/mol |
| $w_{mb}$ | Molecular weight of microbial cells | g/mol |
| $V_g$ | Volume in the gas phase | L |
| $g_1, g_2$ | Polynomial coefficients for volume in the gas phase | |
| $pH$ | Average pH | |
| $R$ | Ideal gas constant | bar·L/(mol·K) |
| **Sigmoid parameters** |  |  |
| $p_1, p_2$ * | Effect inhibition factor of the methanogens growth rate by the action of bromoform | |
| $p_3, p_4$ * | Effect hydrogen control factor for sugar utilization | |
| $p_5, p_6$ * | Effect molar fraction of the sugars utilized via reaction 1 | |
| $p_7, p_8, p_9$ * | Effect molar fraction of the sugars utilized via reaction 2 | |

**Table 2:** Parameter and initial condition values. (Asterisk indicates estimated values, carrot indicates values from virtual microbiome experiments)

| Parameter | Value |  | Initial Value for Estimation |  | Bounds |
| --- | --- | --- | --- | --- | --- |
|  | Control | Treatment | Control | Treatment |  |
| Polymer components (initial conditions) |  |  |  |  |  |
| initial $z_{ndf}$ * | 6.13 | 5.13 | 5.11 | 5.64 | 80 % - 120 % |
| initial $z_{nsc}$ * | 2.02 | 1.86 | 1.68 | 1.68 | 80 % - 120 % |
| initial $z_{pro}$ * | 3.14 | 2.97 | 2.67 | 2.79 | 80 % - 120 % |
| Soluble components (initial conditions) |  |  |  |  |  |
| initial $s_{su}$ | 0 | 0 | | | |
| initial $s_{aa}$ | 0 | 0 | | | |
| initial $s_{ac}$ | 0 | 0 | | | |
| initial $s_{bu}$ | 0 | 0 | | | |
| initial $s_{pr}$ | 0 | 0 | | | |
| initial $s_{IC}$ | 0.14 | 0.14 | | | |
| initial $s_{IN,0}$ | 0 | 0 | | | |
| initial $s_{H_2,0}$ | 0 | 0 | | | |
| initial $s_{CH_4,0}$ | 0 | 0 | | | |
| initial $s_{br,0}$ * | 0 | $3.74 \cdot 10^{-3}$ | | $3.12 \cdot 10^{-3}$ | 80 % - 120 % |
| initial $V_0$ | 0.75 | 0.75 | | | |
| **Gas components (initial conditions)** |  |  |  |  |  |
| initial $n_{g,H_2,0}$ | 0 | 0 | | | |
| initial $n_{g,CO_2,0}$ | 0 | 0 | | | |
| initial $n_{g,CH_4,0}$ | 0 | 0 | | | |
| **Microbial functional groups (parameters)** |  |  |  |  |  |
| $p_{su,0}$ ^ | $2.39 \cdot 10^{-2}$ | $2.667 \cdot 10^{-2}$ | | | |
| $p_{aa,0}$ ^ | $1.0307 \cdot 10^{-1}$ | $8.706 \cdot 10^{-2}$ | | | |
| $p_{m,0}$ ^ | $2.297 \cdot 10^{-2}$ | $2.232 \cdot 10^{-2}$ | | | |
| $TMC$ * | $3.37 \times 10^{-2}$ | $3.37 \times 10^{-2}$ | $6.53 \cdot 10^{-2}$ | $6.53 \cdot 10^{-2}$ | 20 % - 200 % |
| **Rates (parameters)** |  |  |  |  |  |
| $\rho_b$ | $2.34 \times 10^{-2}$ | $2.34 \times 10^{-2}$ | | | |
| $k_{br}$ * | N/A | $1.68 \cdot 10^{-3}$ | N/A | $8.3 \cdot 10^{-3}$ | 20 % - 200 % |
| **Biochemical parameters** |  |  |  |  |  |
| $\sigma_{ch,x}$ | $2.0 \cdot 10^{-1}$ | $2.0 \cdot 10^{-1}$ | | | |
| $\sigma_{pro,x}$ | $5.5 \cdot 10^{-1}$ | $5.5 \cdot 10^{-1}$ | | | |
| $\sigma_{ac,aa}$ | $6.7 \cdot 10^{-1}$ | $6.7 \cdot 10^{-1}$ | | | |
| $\sigma_{pr,aa}$ | $6.2 \cdot 10^{-2}$ | $6.2 \cdot 10^{-2}$ | | | |
| $\sigma_{bu,aa}$ | $2.4 \cdot 10^{-1}$ | $2.4 \cdot 10^{-1}$ | | | |
| $\sigma_{H_2,aa}$ | $8.2 \cdot 10^{-1}$ | $8.2 \cdot 10^{-1}$ | | | |
| $\sigma_{IC,aa}$ | $8.8 \cdot 10^{-1}$ | $8.8 \cdot 10^{-1}$ | | | |
| $k_d$ | $8.33 \cdot 10^{-4}$ | $8.33 \cdot 10^{-4}$ | | | |
| $k_{hyd,ndf}$ * | $4.09 \cdot 10^{-2}$ | $1.24 \cdot 10^{-2}$ | $2.4 \cdot 10^{-2}$ | $2.4 \cdot 10^{-2}$ | 20 % - 200 % |
| $k_{hyd,nsc}$ * | $1.46 \cdot 10^{-1}$ | $1.09 \cdot 10^{-1}$ | $7.38 \cdot 10^{-2}$ | $7.38 \cdot 10^{-2}$ | 20 % - 200 % |
| $k_{hyd,pro}$ * | $1.66 \cdot 10^{-1}$ | $1.59 \cdot 10^{-1}$ | $9.13 \cdot 10^{-2}$ | $9.13 \cdot 10^{-2}$ | 20 % - 200 % |
| $k_{m,aa}$ * | 2.58 | 1.33 | 2.0 | 2.0 | 20 % - 200 % |
| $k_{m,H_2}$ * | 16.26 | 6.84 | 16 | 16 | 20 % - 200 % |
| $k_{m,su}$ * | 2.00 | 1.32 | 1.0 | 1.0 | 20 % - 200 % |
| $K_{S,aa}$ | $6.4 \cdot 10^{-3}$ | $6.4 \cdot 10^{-3}$ | | | |
| $K_{S,H_2}$ | $5.84 \cdot 10^{-6}$ | $5.84 \cdot 10^{-6}$ | | | |
| $K_{S,su}$ | $9.0 \cdot 10^{-3}$ | $9.0 \cdot 10^{-3}$ | | | |
| $K_{S,IN}$ | $2.0 \cdot 10^{-4}$ | $2.0 \cdot 10^{-4}$ | | | |
| $Y_{aa}$ * | $1.61 \cdot 10^{-2}$ | $1.59 \cdot 10^{-1}$ | $3.1 \cdot 10^{-1}$ | $3.1 \cdot 10^{-1}$ | 20 % - 200 % |
| $Y_m$ * | $3.87 \cdot 10^{-3}$ | $1.32 \cdot 10^{-3}$ | $6 \cdot 10^{-3}$ | $6 \cdot 10^{-3}$ | 20 % - 200 % |
| $Y_{su}$ * | $1.04 \cdot 10^{-1}$ | $4.81 \cdot 10^{-2}$ | $1.6 \cdot 10^{-1}$ | $1.6 \cdot 10^{-1}$ | 20 % - 200 % |
| **Physiochemical parameters** |  |  |  |  |  |
| $K_{a,CO_2}$ | $5.03 \cdot 10^{-7}$ | $5.03 \cdot 10^{-7}$ | | | |
| $kLa_{CH_4}$ * | 4.58 | 8.41 | 4.28 | 4.28 | 20 % - 200 % |
| $kLa_{CO_2}$ * | 4.07 | 8.25 | 4.28 | 4.28 | 20 % - 200 % |
| $kLa_{H_2}$ * | 4.40 | 8.35 | 4.28 | 4.28 | 20 % - 200 % |
| $K_{H,CO_2}$ | $2.59 \cdot 10^{-2}$ | $2.59 \cdot 10^{-2}$ | | | |
| $K_{H,CH_4}$ | $0.1 \cdot 10^{-3}$ | $0.1 \cdot 10^{-3}$ | | | |
| $K_{H,H_2}$ | $7.31 \cdot 10^{-4}$ | $7.31 \cdot 10^{-4}$ | | | |
| $T$ | 310.15 | 310.15 | | | |
| $t_{full}$ | 10.68 | 10.68 | | | |
| $w_{su}$ | 180.16 | 180.16 | | | |
| $w_{aa}$ | 134 | 134 | | | |
| $w_{mb}$ | 113 | 113 | | | |
| $g_1$ | $1.59 \cdot 10^{-2}$ | $7.99 \cdot 10^{-3}$ | | | |
| $g_2$ | $5.48 \cdot 10^{-2}$ | $3.60 \cdot 10^{-2}$ | | | |
| $pH$ | 6.8 | 6.8 | | | |
| $R$ | $8.314 \cdot 10^{-2}$ | $8.314 \cdot 10^{-2}$ | | | |
| **Sigmoid parameters** |  |  |  |  |  |
| $p_1^*$ | 1694.6 | 1694.8 | 1694.6 | 1694.6 | 20 % - 200 % |
| $p_2^*$ | $2.14 \cdot 10^{-4}$ | $2.62 \times 10^{-5}$ | $1.08 \cdot 10^{-4}$ | $1.08 \cdot 10^{-4}$ | 20 % - 200 % |
| $p_3$ | 102.5300 | 102.5300 | | | |
| $p_4$ | $2.35 \cdot 10^{-1}$ | $2.35 \cdot 10^{-1}$ | | | |
| $p_5$ | 7.5 | 7.5 | | | |
| $p_6$ | $6.98 \cdot 10^{-2}$ | $6.98 \cdot 10^{-2}$ | | | |
| $p_7$ | $9.219 \cdot 10^{-1}$ | $9.219 \cdot 10^{-1}$ | | | |
| $p_8$ | 2.55 | 2.55 | | | |
| $p_9$ | $1.68 \cdot 10^{-1}$ | $1.68 \cdot 10^{-1}$ | | | |

### 2.3 Parameter Estimation

Muñoz-Tamayo et al. (12) estimated parameters using the experimental data from Chagas et al. (24) and the original model, which did not account for *A. taxiformis’* effect on rumen fermentation, Muñoz-Tamayo et al. (13), estimated parameters using the experimental data from Serment et al. (25). Muñoz-Tamayo et al (12) estimated parameters with the first 24 hours of Chagas et al. (24) data of *A. taxiformis* at five different inclusion rates and a negative control in a serum bottle system. Data used in the estimation included acetate, butyrate, propionate, NH_3_, and the moles of methane produced. Muñoz-Tamayo et al. (13) estimated parameters with data from four treatment combinations of different inoculum and substrate types from a 24-hour syringe system in Serment et al. (25). Data used in the estimation included acetate, butyrate, propionate, inorganic nitrogen, pH, and moles of CH_4_ and CO_2_.

The experimental set ups of the *in vitro* studies used to calibrate the Muñoz-Tamayo et al. (13) and Muñoz-Tamayo et al. (12) models have significant differences with the Roque et al. (3) RUSITEC *in vitro* study. Due to the different *in vitro* systems used, parameters estimated in Muñoz-Tamayo et al.(13)and Muñoz-Tamayo et al. (12) parameters are not directly comparable, and parameters were estimated using Roque et al. (3) data.

Model parameters were estimated with MATLAB’s optimization toolbox constrained nonlinear optimization solver **fmincon** with default values for termination tolerances. Parameters were varied over prescribed bounds to minimize the cost function of the sum of the squared differences between model CH_4_ and CO_2_ output and CH_4_ and CO_2_ data for 96 hours for control and treatment conditions.

Parameters that were estimated, their prescribed bounds, and nominal values can be found in Table 2. Estimated parameters include 17 parameters varied separately for control and treatment conditions, two parameters estimated only in treatment conditions (the kinetic rate constant of bromoform utilization, and the initial bromoform concentration), and one parameter estimated together for control and treatment conditions (total microbial concentration). We are assuming that both the control and treatment responses were generated by a different parameter set given that they have different initial conditions and feed compositions.

### 2.4 Classification Methodology

Recognition of a distinctive microbial profile specific to individual ruminants was first observed within the protozoal population (26,27) and then in the fibrolytic bacterial community (28). Since the advent of more advanced molecular sequencing tools, it is widely accepted that the cattle rumen has a common set of associated (29), but has large variation outside of it (30)

Muñoz-Tamayo et al. (12) considered a macroscopic representation of fermentation: the microbiota was aggregated into three functional groups with fixed chemical compositions defined by the monomers included in the model that they use as substrates. Other models have often used a single microbial group or two pools of microbes (5,6). While including only a limited number of functional groups is a clear simplification of the microbiome, Muñoz-Tamayo et al. (12)’s approach gives a good prediction of rumen fermentation.

Due to the importance of the microbial community in rumen fermentation and the variation in microbial populations in both functionality and abundance, we used phylogenetic data to determine the initial conditions of the three functional microbial groups (sugar utilizers, amino acid utilizers, and methanogens). The functionality of the microbes from the phylogenetic data had to be assigned to determine these initial conditions. We identified key species based on Kettle et al. (31) functional classifications.

Phylogenetic microbial data in Roque et al. (3) yields the relative abundance of the microbes present in the *in vitro* experiment. This data is normalized and fails to provide an absolute quantification of taxa. Thus, the relative abundance microbial data was then converted to microbial group concentrations by normalizing the quantities by the total microbial concentration (TMC (mol/L)).

TMC was estimated during parameter estimation (initial guess was from Nagadi et al. (32) *in vitro* experiment value). To classify the microbes identified in Roque et al. (3) based on functionality, specifics from the functional groups defined by Kettle et al. (31) were used to determine search terms to assign the data from Roque et al. (3) into the three functional microbial groups. Between 82% and 92% of microbes across all time points were functionally unclassified using this classification procedure. To increase the accuracy of model predictions, further classification was necessary.

This preliminary classification divided the unclassified microbes into four sets: relative abundance of functionally classified amino acid utilizers, sugar utilizers, and methanogens, with unknown genus level (*u*_*aa*_, *u*_*su*_,and *u*_*u*_ respectively) and relative abundance of functionally unclassified (*u*_*u*_).

Due to the uncertainty of classification of functionally classified microbes with unknown genus level and the large number of functionally unclassified microbes, the classification of microbes was further investigated. Variation was added to the above-described data set by randomly assigning functionally unclassified microbes and microbes with unclassified genus level to the different functional groups. It is not computationally feasible to test every combination of microbe in each functional group, as it would take more than 10^768^ model simulations. Thus, we performed simulations with random initial conditions of the three functional microbial groups.

*u*_*aa*_, *u*_*su*_, *u*_*m*_, *and u*_*u*_,were randomly partitioned into relative abundance of amino acid utilizers (*p*_*aa*_), relative abundance of sugar utilizers (*p*_*aa*_), relative abundance of methanogens (*p*_*m*_), and relative abundance of other utilizers (*p*_*ou*_). To randomly partition the four sets of unclassified microbes, 16 random distribution parameters were varied simultaneously and uniformly. The distribution parameters determined the fractions of unclassified microbes that were partitioned into different functional groups. The distribution parameters are denoted as *D*_*i,j*_, where i are the unclassified microbes and j is the functional group to which they are partitioned. The relative abundance of the functional microbial group *i* can be written as

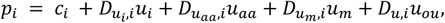

where *c*_*i*_ denotes the relative abundance of functionally classified microbes with known genus, computed using the initial classification procedure.

We explored our parameter space of *D*_*ij*_’s using Satlelli’s extension of the Sobol’ sequence (33), using a Pythob sensitivity analysis library (SALib) (34,35). The Sobol sequence is a quasi-random low-discrepancy sequence, meaning it is a deterministic sequence that converges quickly to a uniform distribution and fills the space of possibilities evenly. We generated N sets of 16 floats on (0,1) using the Sobol’ sequence. To ensure that the total microbial population stayed constant across sampling, it was necessary to normalize the floats so that 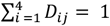 for all j.

Saletlli’s extension of Sobol’s sequence reduces error rates in the sensitivity indices and requires fewer samples (33). For the Sobol sequence to converge to a uniform distribution, *N* = *n*(16 + 2) samples are needed, with 16 the number of uncertain inputs, and n the number of samples. We used n = 16384 for calculations of both control and treatment virtual microbiome experiments.

### 2.5 Sensitivity Analysis

Next, we detail the mathematical and statistical methods used in our local and global sensitivity analyses.

#### 2.5.1 Global Sensitivity Analysis

Global sensitivity analysis (GSA) considers the impact of varying parameters simultaneously and uniformly over a range of possible values, quantifies model sensitivity by the variance of model output, and attributes fractions of output variance to model inputs (36). In contrast to local sensitivity analysis (LSA) methods, GSA methods are helpful when models are nonlinear and nonadditive and have a high degree of dimensionality. We estimate the effects of parameter variations by using Satlelli’s extension of the Sobol’ sequence (33) using Python’s sensitivity analysis library SALib. (34,35) The Sobol sequence generated N sets of M parameters within prescribed bounds. We quantified the variance of model output using Sobol sensitivity indices (33,36).

Sobol sensitivity indices decompose variation of model output and attribute these changes to individual parameters or the interactions between multi parameter changes. Final methane production was the model output that was considered in the analysis. Assuming that parameters are independent, the variance of final methane production, m, with respect to random parameter inputs *x* = (*x*_1_, …, *x*_*k*_), may be decomposed as

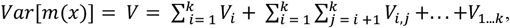

where *V* is the total variance, *V*_*i*_ is the first order effects of parameter *i*, and *V*_1,…*n*_ is the contribution to the variance from 1^st^, 2^nd^,…, and n^th^ parameter. The total variance is the sum of first order effects of the parameters as well as higher order effects from interactions between multi parameter changes. The first order Sobol index of *i* (*S*_*i*_) is defined as the contribution of parameter *i* to total variance:

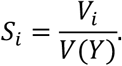

*S*_*i*_ is normalized since: 0 ≤ *V*_*i*_ ≤ *Var*(*Y*). s

The total-order Sobol index of 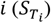 captures the effect on total variance of parameter *i*. It includes the contribution of parameter *i* to total variance as well as all interaction terms that include parameter *i*. It can be written as:

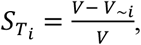

where *V*_∼*i*_ is the sum over the set of all variance terms not containing the *i*_th_ parameter. When the model is only additive, 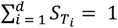.

We used a Python sensitivity analysis library (SALib) for the estimation of sensitivity indices. (34,35) *N* = *n*(*M* + 2) model evaluations were needed, with M the number of uncertain inputs, and n the number of samples. We used n = 16384 for all calculations, as this was sufficient for convergence of the first and total order Sobol indices.

#### 2.5.2 Local Sensitivity Analysis

Local sensitivity analysis methods quantify the variation in model output when parameters are perturbed one at a time (OAT). These methods do not consider parameters varying in combination with each other. LSAs are helpful when model inputs behave linearly or additively (36). Prescribed parameters were varied between prescribed bounds one at a time and final methane model output was measured each time.

## 3 Results

### 3.1 Parameter Estimation

Figure 2 shows the model fits of CH_4_ and CO_2_ concentrations from RUSITEC samples generated with and without *A. taxiformis* added to the feed. The parameters that were held constant and the parameters that were estimated can be found in Table 2.

**Figure 2:**
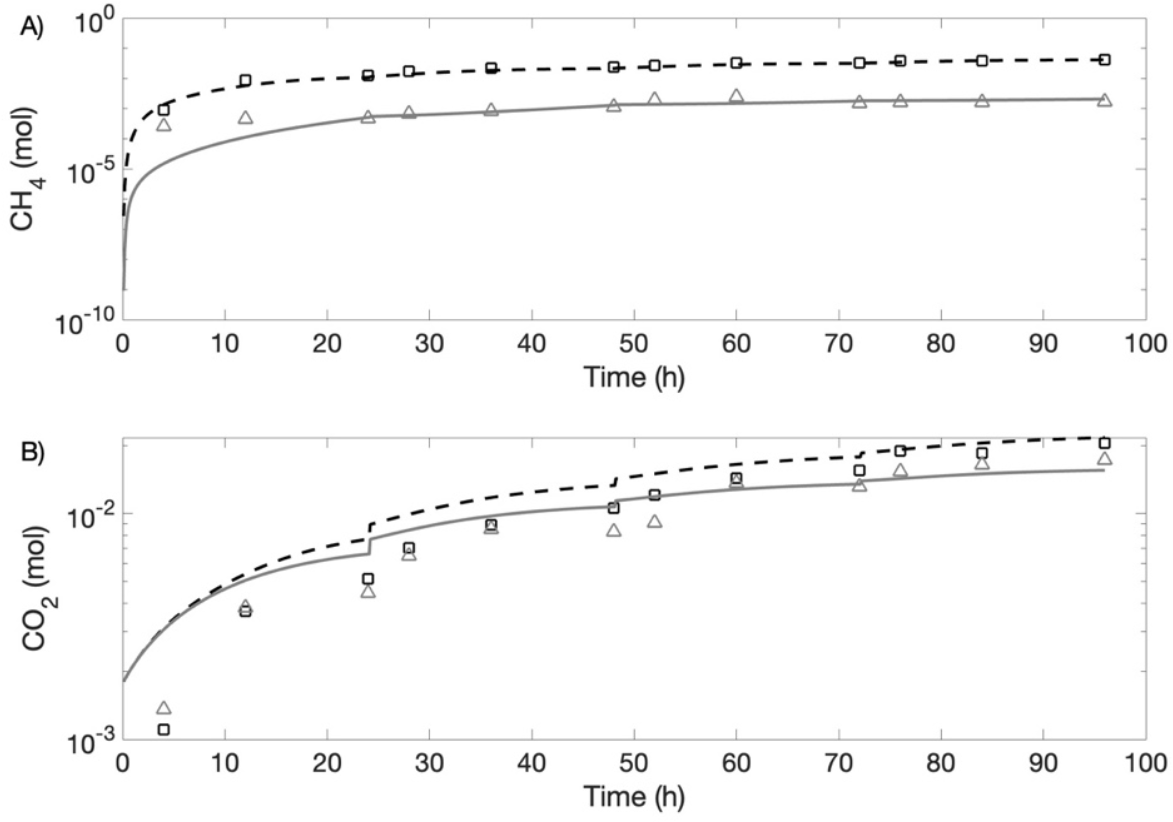
Model calibration with experimental data after initial parameter estimation. Experimental data of treatment (square) and control (triangle) are compared against model response of treatment (**- - -**) and control (**-—**) for (A,B) CH_4_ and (C,D) CO_2_.

To quantitatively assess the accuracy of the fits, we performed a linear fit of prediction as a function of observation, to describe the relationship between the data and model output. The linear fitted curve was calculated using the least squares method, by minimizing the sum of the square of errors between the data and the model output. The smaller the absolute value of the y-intercept and the closer the slope is to one, the better the model prediction. The observed data plotted against the prediction of the fits, as well as the calculated best fit line CH_4_ and CO_2_ are shown in Figure 3. The linear regression equations of control and treatment CH_4_ and control and treatment CO_2_ had slopes of 0.99, 0.80, 1.05, and 1.27 respectively and all had y-intercepts with absolute value than 4 · 10^−3^. This shows that the model satisfactorily captures the dynamical behavior of both CH_4_ and CO_2_.

**Figure 3:**
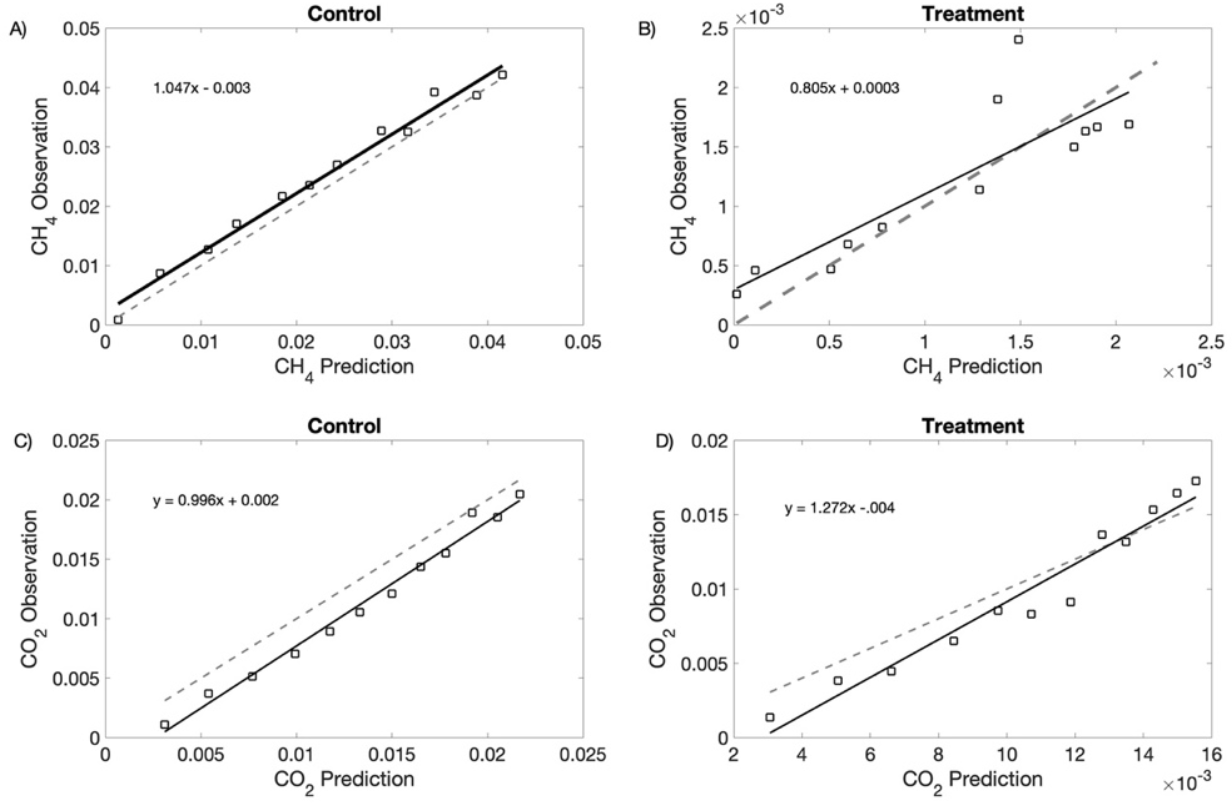
Observations plotted against model predictions (square) before virtual microbiome experiments. Linear fitted curve is the solid black line (equation of line displayed on graph). Line y = x is the gray dashed line. The linearity shows the reliability of the model.

### 3.2 Virtual Microbiome Experiments

After the classification of microbial relative abundance data, between 82% and 92% of microbes remained functionally unclassified across all time points. The variation added to this data set, as described in the classification methodology, produced additional sets of data with different distributions of functionally unclassified and unclassified at the genus level microbes. Each data set was used to compute the initial conditions of the functional groups for a model simulation.

The methane output of the virtual microbiome experiments can be seen in Figure 4. The spread in control compared to in treatment shows that the addition of *A. taxiformis* makes the system more sensitive to microbial variation. The simulations that produced the final methane closest to Roque et al. (3) data are indicated with a red curve, the 1st percentiles are indicated in blue, and the 0.1st percentiles are indicated in cyan on the graphs. In the treatment case, most model simulations resulted in methane output substantially larger than Roque et al. (3), and the simulation that produced the final methane closest to Roque et al. (3) data (red curve on graph) fell below the 1st percentile of methane output. The large variations demonstrate the importance of functional classification of microorganisms for accurate methane prediction. In the control case, most model simulations were close to or lower than Roque et al. (3) data. This suggests that the system is more sensitive with bromoform.

**Figure 4:**
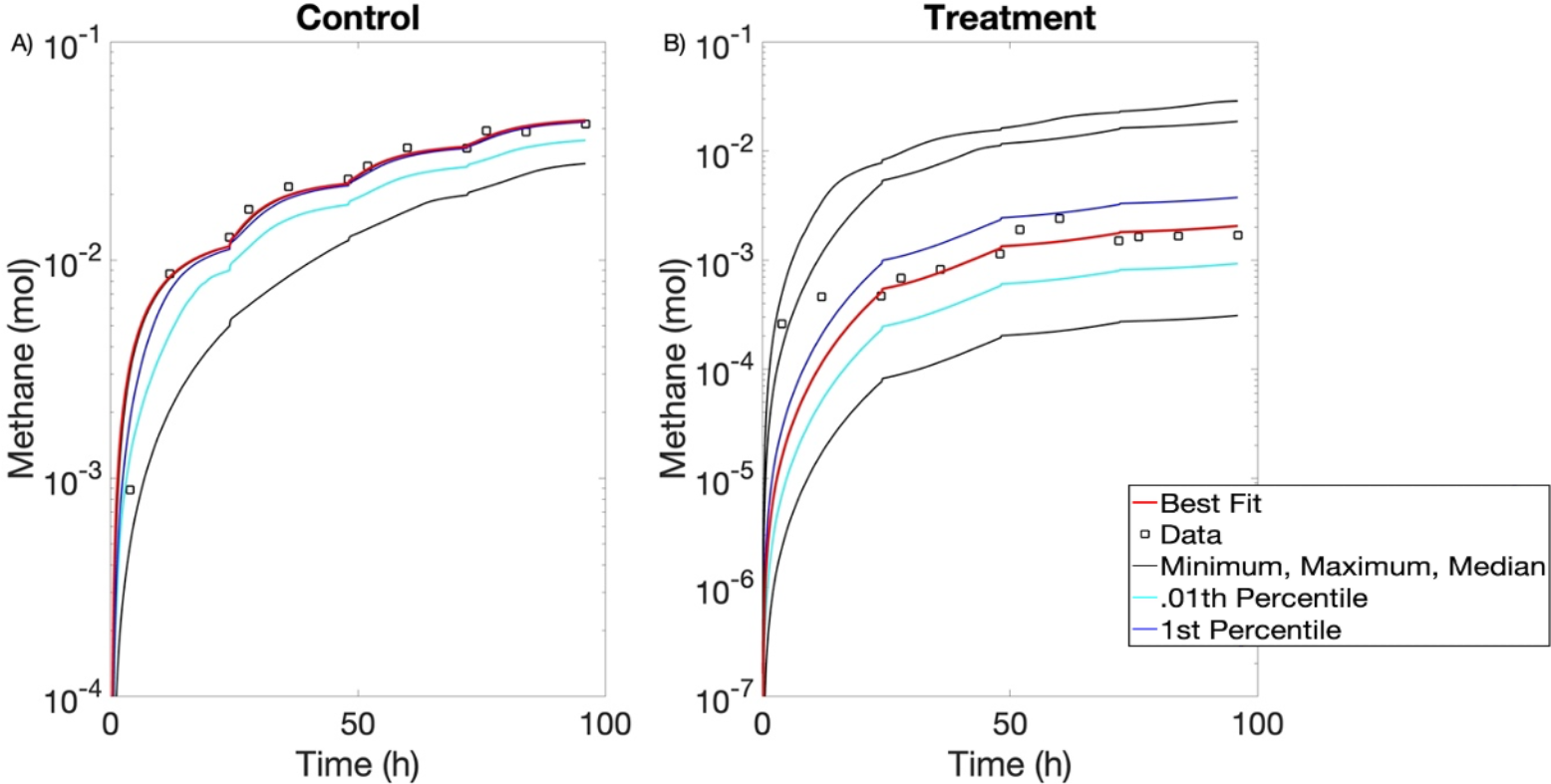
Methane output of virtual microbiome experiments. Experimental data (square) compared against model output of varying distributions of initial values of functional microbial group concentrations. Red line corresponds to the simulation that best fits the data. Max, min, and median indicated in black.1st percentile shown in blue and .01st percentile shown in cyan. (A) Control. (B) Treatment.

Figure 5 includes histograms of the initial values for relative abundances of functional microbial groups used in the virtual microbiome experiments. The relative abundances used to calculate the initial values of the functional microbial groups that produced the model simulation with final methane closest to Roque et al. (3) data are indicated with dashed black lines (same simulation as indicated with the red curves in Figure 4). Table 3 includes the resulting concentrations of functional microbial groups. The virtual microbiome experiments predict that bromoform has effects on the microbiome diversity, even beyond the reduction in methanogens. The virtual microbiome experiments placed 73.6% into sugar utilizers, 15.1% into amino acid utilizers, and 6.8% into methanogens in control conditions and 31.8% into sugar utilizers, 58.5% into amino acid utilizers, and 2.2% into methanogens in treatment conditions, as seen in Figure 5. The virtual microbiome experiments for control and treatment conditions placed 4.5% and 7.5% respectively of the virtual microbiome into a functionally unclassified category. This suggests that including additional functional groups might be needed for a more complete representation of the rumen microbiome. Additionally, Figure 5 shows that in both control and treatment conditions, the best methane fit is produced by relative abundances of methanogens on the left tail of the distribution of methanogens tested. This means very few methanogens produce the best methane fit, which suggests that most of the functionally unclassified microbes are not methanogens in either condition.

**Table 3:**
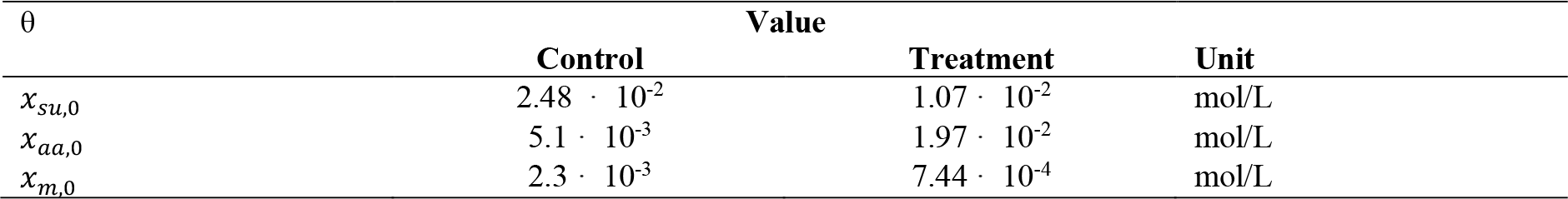
Initial Conditions from Virtual Microbiome.

| $\theta$ | Value | | |
| --- | --- | --- | --- |
|  | Control | Treatment | Unit |
| $x_{su,0}$ | $2.48 \cdot 10^{-2}$ | $1.07 \cdot 10^{-2}$ | mol/L |
| $x_{aa,0}$ | $5.1 \cdot 10^{-3}$ | $1.97 \cdot 10^{-2}$ | mol/L |
| $x_{m,0}$ | $2.3 \cdot 10^{-3}$ | $7.44 \cdot 10^{-4}$ | mol/L |

**Figure 5:**
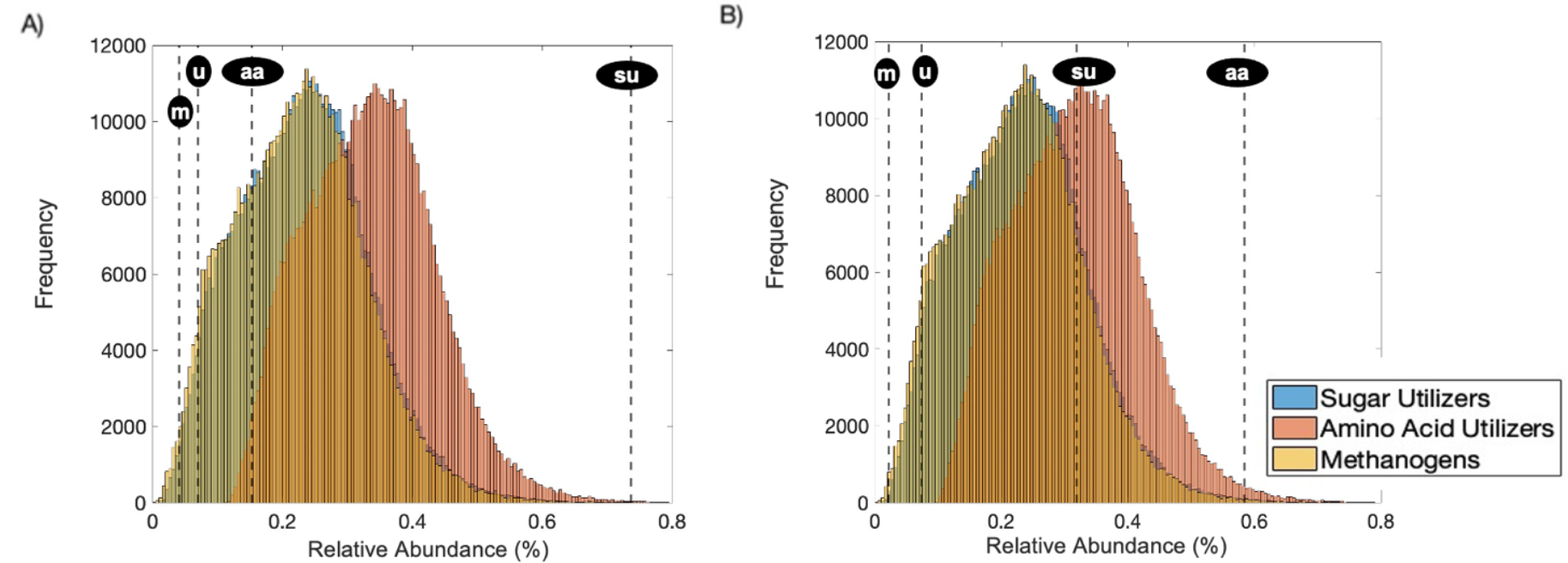
Relative abundances of virtual microbiome experiments. Histograms of initial values for relative abundances of functional microbial groups used in virtual microbiome experiments. Black dashed lines (- - -) correspond to initial values of relative abundances of functional microbial groups that result in best methane fit (unclassified (u), methanogens (m), amino acid utilizers (aa), sugar utilizers (su)). (A) Control. (B) Treatment.

Figure 6 shows the CH_4_ and CO_2_ data from Roque et al. (3) with and without *A. taxiformis* compared against the model’s prediction with parameter values from the initial parameter estimation and microbial functional group initial conditions that produced the best fit identified in the virtual microbiome experiments. To provide an indication of the reliability of the model, the linear fitted curve was calculated which is shown in figure 7. The linear fitted curve was calculated using the least squares method, by minimizing the sum of the square of errors between the data and the model output and describes the relationship between the data and model output. The smaller the absolute value of the y-intercept and the closer the slope to one, the better the model prediction. The model satisfactorily captures the behavior of both CO_2_ and CH_4_. The linear regression equations of control and treatment CH_4_ and control and treatment CO_2_ had slopes of 0.99, 0.81, 1.05, and 1.26 respectively and all had y-intercepts with absolute value than 4 · 10^−3^. These fits are not quantitatively better than before the virtual microbiome experiments since the regression equations have similar slopes and y-intercepts. However, they include a fuller representation of the rumen microbiome, whose function is vital in rumen fermentation. Going forward the parameter set after the virtual microbiome experiments is used since it includes a greater proportion of microbes and is still able to satisfactory predict CO_2_ and CH_4_ output.

**Figure 6:**
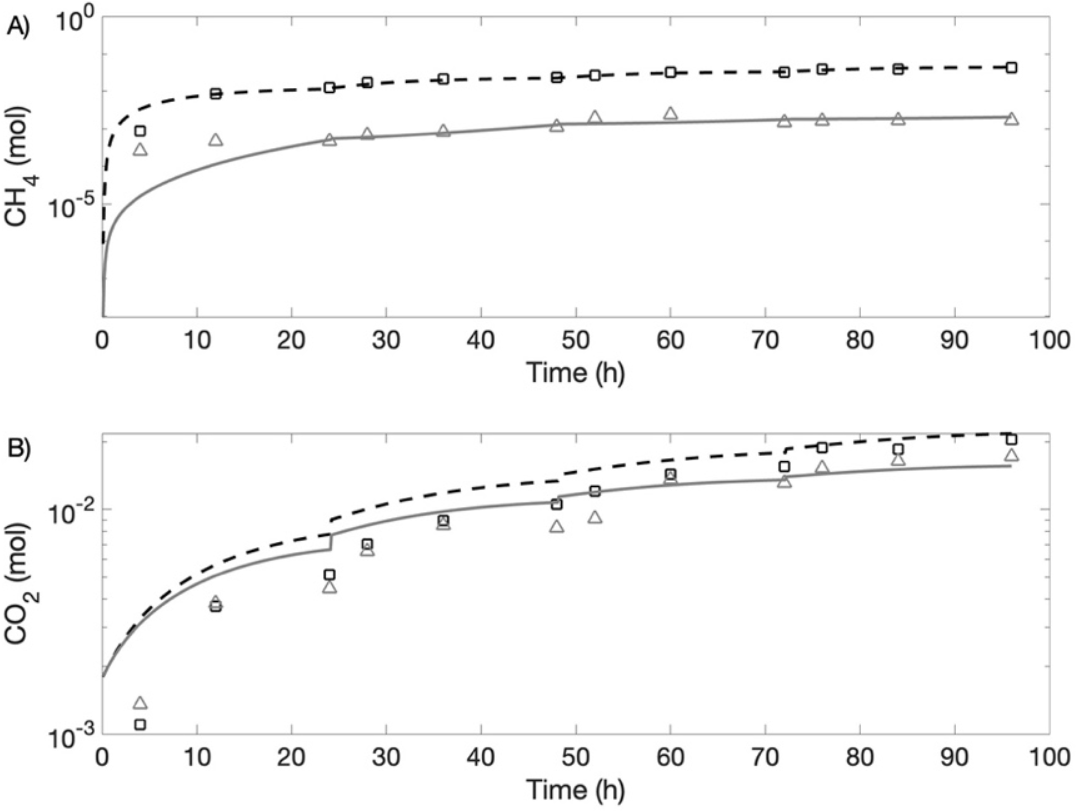
Model calibration with experimental data after Virtual Microbiome Experiments. Experimental data of treatment (square) and control (triangle) are compared against model response of treatment (**- - -**) and control (**-—**) for CH_4_ and CO_2_.

**Figure 7:**
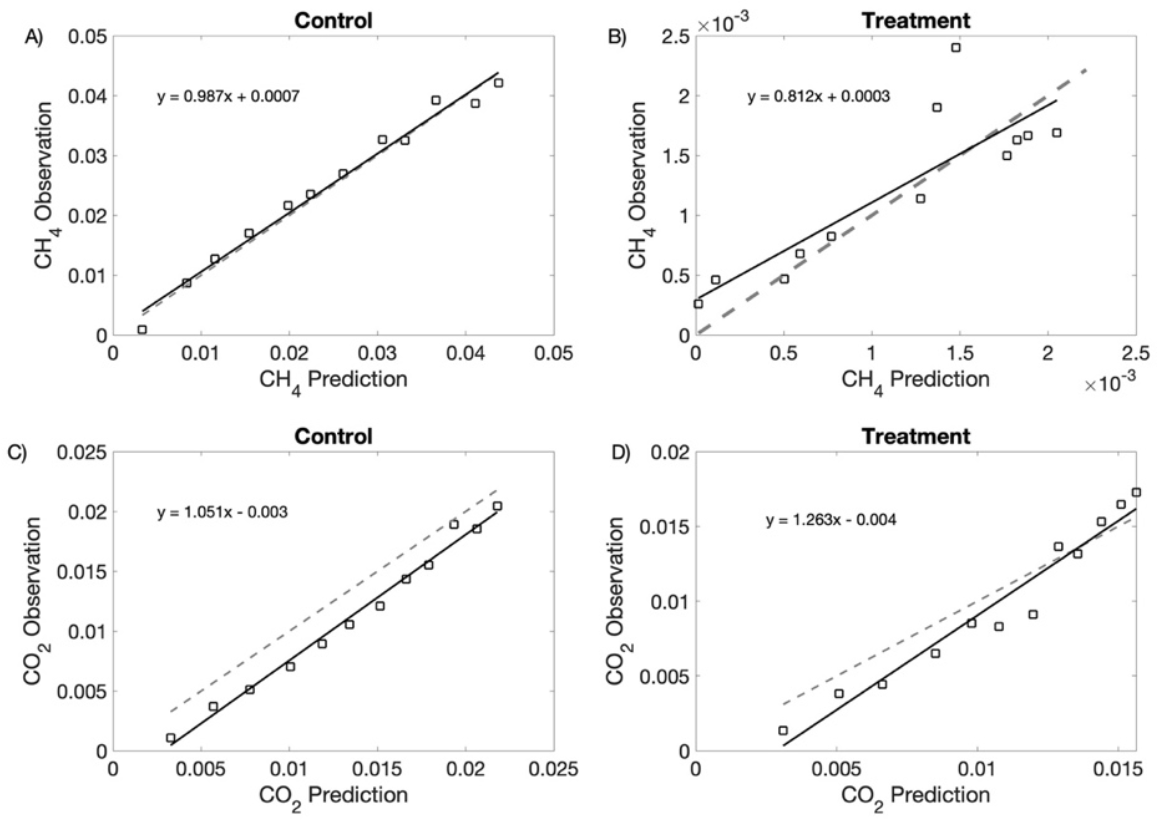
Observations plotted against model predictions (square). Linear fitted curve is the solid black line (equation of line displayed on graph). Line y = x is the gray dashed line. The linearity shows the reliability of the model.

### 3.3 Sensitivity Analysis

In this section, a sensitivity analysis is performed on all model parameters without a significant source in literature or a certain experimental value (parameters indicated with a * in Table 1).

We examined the sensitivity of the model’s output of the final gaseous methane for two conditions:

1. Control: Roque et al. (3) control experimental conditions and parameter values that were estimated for the control conditions.
2. Treatment: Roque et al. (3) treatment experimental conditions and parameter values that were estimated for treatment conditions.

A global sensitivity analysis was performed for all conditions. Parameters were varied simultaneously between 50-150% of their baseline values. The model was evaluated 540,672 times for control and 573,440 times for treatment conditions.

The coefficient of variation in the final methane output is 0.23 and 1.22 for control and treatment respectively. The coefficient of variation is the standard deviation divided by the mean and is provided as a scale for the Sobol indices between conditions. It is significantly higher for treatment, which indicates, similarly to microbial classification, that the final methane output showed larger variation when bromoform is present.

The results of the global sensitivity analysis for each parameter whose total order Sobol index was greater than 0.05 for control or treatment conditions for control and treatment can be found in Figure 8 AB.

**Figure 8:**
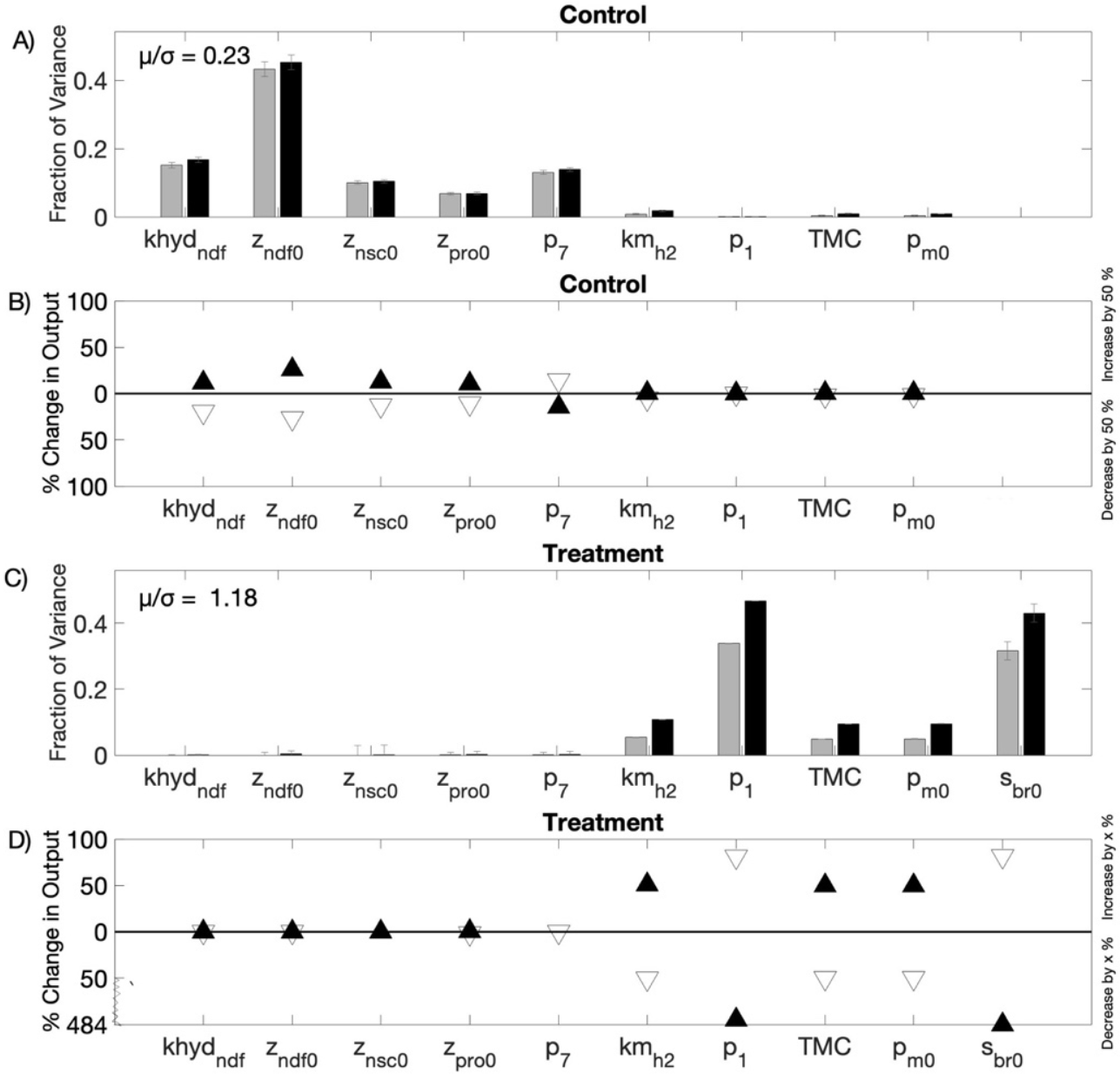
Sensitivity Analysis. *Global Sensitivity Analysis (A,C)*: First (gray) and Total (black) Order Sobol indices are plotted as bars with errors of 95% confidence interval. The coefficient of variation (*μ/σ*) is displayed, for comparing the fraction of variance between conditions. *Local Sensitivity Analysis (B,D):* Triangles represent percentage change of final gaseous methane from baseline model output for parameters. The filled upward or outlined downward triangles indicate the direction of variation. Parameters were quantified by the absolute difference they produced when they were decreased and increased by x = 50% for all parameters.

The total order Sobol index is greater than 0.05 for five control and five treatment parameters. The selected parameter sets do not include any common parameters. The control included the three feed initial conditions (z_ndf0_, z_nsc0_, z_pro0_), one hydrolysis rate constant (khyd_ndf_), a parameter affecting the maximum of a sigmoid function of the fraction of sugar utilization via reaction 2 (p_7_; see Supplement A for details). Treatment included the total microbial concentration (TMC), the initial percent abundance of methanogens (p_m0_), the maximum specific utilization rate of hydrogen (km_h2_), a parameter affecting the steepness of a sigmoid function of the inhibition factor of the methanogens growth rate by the action of bromoform (p_1_; see Supplement A for details), and initial bromoform concentration (s_br0_).

For the control condition, the initial neutral detergent fiber concentration (z_nd0_) has the largest individual effect in the global sensitivity analysis (see Figure 8A). It accounts for 43% of the total model variance in this condition on its own. The initial nonstructural carbohydrate concentration (z_scf0_), the initial protein concentration (z_pro0_), the hydrolysis rate of cell wall carbohydrates (khyd_ndf_), and a parameter affecting thee fraction of sugar utilization via reaction 2 (p_7_) all contribute between 6-16% of total model variance individually, while the total microbial concentration (TMC), the initial percent abundance of methanogens (p_m0_), the maximum specific utilization rate of hydrogen (km_h2_), a parameter that effects the inhibition factor of methanogens growth rate by the action of bromoform (p_1_), and initial bromoform concentration (s_br0_) have almost zero both first and total order effects. The three feed initial conditions (z_nsc0,_ z_ndf0_, z_pro0_) together account for 60.1% of the total model variance suggesting the importance of feed composition for methane reduction strategies. Additionally, the low Sobol indices for p_m0_ and TMC suggest that small changes in the microbial functional groups do not have large effects on methane production, which supports the findings of the virtual microbiome experiments.

For the treatment parameters the initial bromoform concentration (s_br0_), and a parameter that effects the inhibition factor of methanogens growth rate by the action of bromoform (p_1_) have the largest individual effects (see Figure 8B). They account for 46.0 % and 45.6% of the total model variance on their own respectively. The maximum specific utilization rate of hydrogen (km_h2_), the initial methanogen relative abundance (p_m0_) and the total microbial concentration (TMC) contribute to 10.4%, 9.21% and 9.14% respectively, while the three feed initial conditions (z_ndf0_, z_nsc0_, z_pro0_), a hydrolysis rate constant (khyd_ndf_), and a parameter affecting the fraction of sugar utilization via reaction 2 (p_7_) have almost zero both first and total order effects.

Comparing first order and total order Sobol indices can quantify the significance of parameter interactions. All five parameters whose Sobol index was greater than 0.05 for treatment conditions have first order and total order Sobol indices that are statistically different. This is not the case in control conditions, which means that parameters in treatment conditions have more higher order effects.

When conducting local sensitivity analysis, we found that despite the model being nonlinear, the final methane output (our metric of interest in the sensitivity analysis) does not change or behaves monotonically when any parameter is varied OAT. In Figure 9, we show this behavior for parameters in both control and treatment conditions whose total order Sobol index was greater than 0.05 for control or treatment conditions.

**Figure 9:**
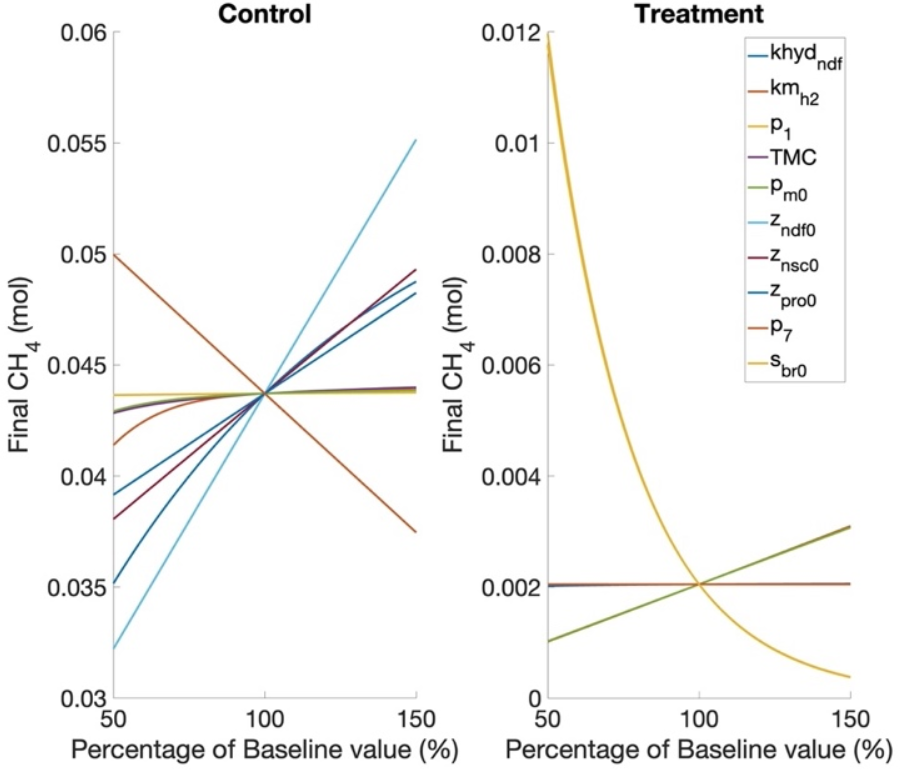
Monotonicity of change in final methane due to variation in parameter values. Parameters whose total order Sobol index was greater than 0.05 for control or treatment are graphed.

The monotonic behavior of parameters when varied OAT allowed us to quantify the sensitivity of model parameters using only their extremal values, using the same method as in Link et al. (37). The local sensitivity analysis used three values: the model was evaluated at 50%, 100%, and 150% of baseline values. The percentage change of final gaseous methane from baseline model output and direction of variation was calculated for each model simulation. Figure 8CD, show the results of the local sensitivity analysis for parameter whose total order Sobol index was greater than 0.05 for control or treatment conditions.

The local sensitivity shows if the parameters are positively or negatively correlated with methane output. In control conditions the three feed initial conditions (z_nsc0,_ z_ndf0_, z_pro0_) and the hydrolysis rate of cell wall carbohydrates (khyd_ndf_), are positively correlated, with a 50% increase in parameter value increasing methane production by 12.8%, 26.2%, 10.4%, and 11.5% repetitively, and a 50% decrease in parameter value decreasing methane production by 12.9%, 26.3%, 10.4%, and 19.6% respectively. This suggests a possibility of methane reduction by changing feed composition, as a reduction of initial neutral detergent fiber concentration (z_nd0_) reduces methane more than an equivalent increase of initial nonstructural carbohydrate concentration (z_scf0_) and the initial protein concentration (z_pro0_) increases methane. Additionally, the local sensitivity analysis shows that a parameter affecting the fraction of sugar utilization via reaction 2 (p_7_) is negatively correlated with a 50% increase in p_7_ decreasing methane production by 11.5% and a 50% decrease in p_7_ increasing methane production by 14.3%.

The local sensitivity analysis for treatment shows that the maximum specific utilization rate of hydrogen (km_h2_), the initial relative abundance of methanogens (p_m0_) and the total microbial concentration (TMC) are positively correlated with a 50% increase in the parameter value increasing methane production by 51.1%, 49.9% and 49.9% respectively and a 50% decrease in parameter value decreasing methane production by 50.4%, 50.0%, and 50.0% respectively. The initial bromoform concentration (s_br0_) and a parameter that effects the inhibition factor of methanogens growth rate by the action of bromoform (p_1_) are negatively correlated with a 50% increase in the parameter value decreasing methane production by 81.17% and 81.4% respectively and a 50% decrease in parameter value increasing methane production by 483.8% and 473.8% respectively.

In the global sensitivity analysis, the sum of all first-order effects accounts for 96.1% and 80.4% of the total variance of control and treatment respectively. This shows that the system is dominated by first-order effects for both conditions and helps explain the similar trends in the global and local sensitivity analysis results. In treatment higher order effects contribute more to overall variance, which suggests that there are some interaction effects due to bromoform in the system.

The control sensitivity analysis suggests that there is possibility of changing methane output by manipulating feed parameters without adding bromoform to the system. It also shows that microbial community variation does not have a significant effect on methane output given the parameter bounds set in the sensitivity analysis in control conditions. However, in treatment conditions microbial community variation has a significant effect on methane output. During the microbe classification, a similar trend was evident: methane output was more sensitive to microbial distributions in treatment conditions compared to control.

We completed an additional sensitivity analysis after the secondary parameter estimation and a similar parameter set was identified (see supplement B).

### 3.4 Further Analysis of Parameters

#### 3.4.1 Control Optimization Guided by Sensitivity Analysis

Further analysis of the system was guided through the results of the sensitivity analysis. Some parameters that were identified as significant in the sensitivity analysis and that are adjustable through feeding adjustments were investigated. In control conditions, final methane output is sensitive to feed parameters including the initial neutral detergent fiber concentration (z_ndf0_), initial nonstructural carbohydrate concentration (z_nsc0_), and the initial protein concentration (z_pro0_). The constrained minimization method was used to find parameter values of the feed initial conditions (z_pro,0_, z_ndf,0_, z_nsc,0_) that minimized methane production at each timepoint in control conditions.

Parameters were allowed to vary 50% from their estimated parameter values. Additionally, the optimization was constrained under the condition that the total substrate hydrolyzed had to remain constant:

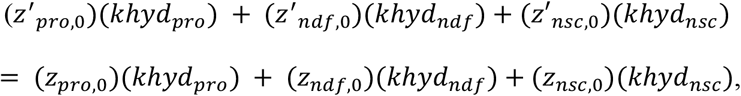

where *z*′ _*i*, 0;_ is the varying feed initial condition i and *z*_*i*, 0_ is the estimated feed initial condition i. These constraints were included to ensure that necessary nutrients were not removed.

Parameter values that minimized methane production can be found in Table 4. This reduction in methane is predicted when initial neutral detergent fiber concentration is reduced by 49.7%, initial nonstructural carbohydrate concentration is reduced by 38.0%, and initial protein concentration is increased by 49.6%. The percent change column shows how all optimized parameter values are close to their bounds and suggests that if parameters are allowed to vary more than 50% from baseline that we could see an even greater reduction in methane output; however, the bounds are necessary to ensure that sufficient nutrients are provided to the animal.

**Table 4:** Optimization parameter values.

| Parameter | Before Optimization | After Optimization | % Change |
| --- | --- | --- | --- |
| $z_{ndf}$ | 6.13 | 3.08 | -49.7% |
| $z_{nsc}$ | 2.02 | 1.25 | -38.0% |
| $z_{pro}$ | 3.14 | 4.69 | +49.6% |

The optimization problem identified parameter values that resulted in a 25.6% reduction of methane from baseline control parameters. This reduction of methane compared to baseline control parameters can be seen in Figure 10.

**Figure 10:**
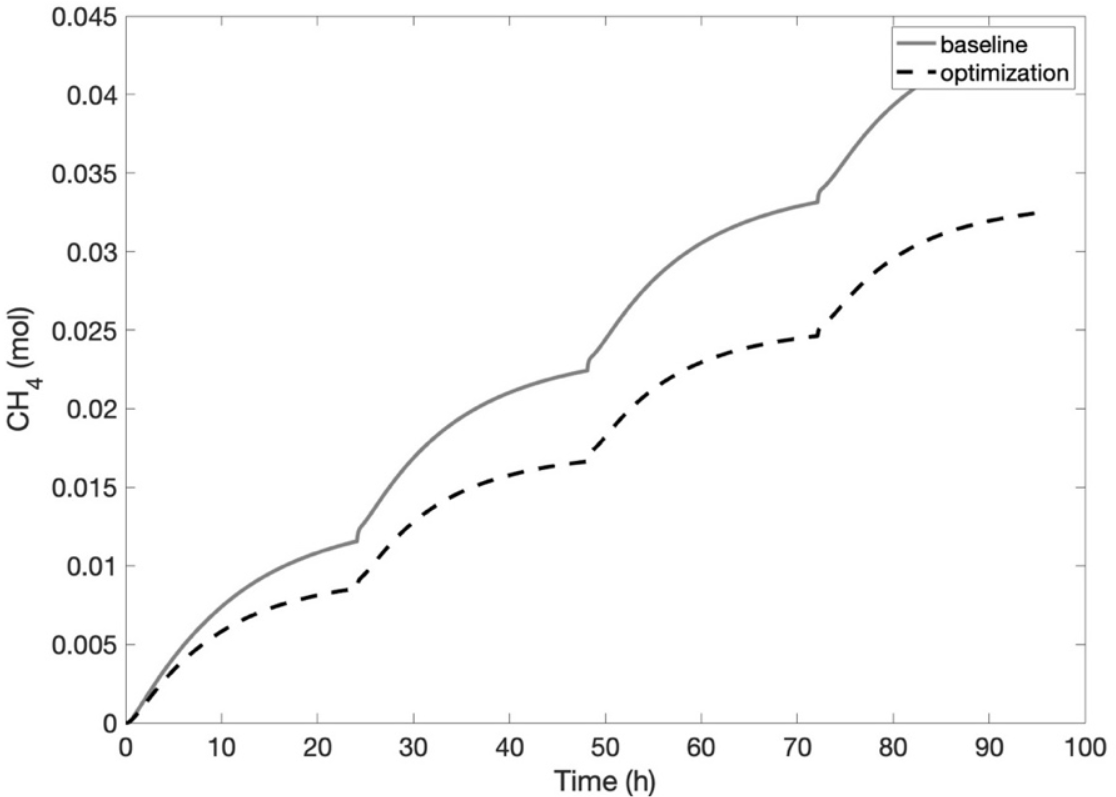
Methane output of optimization (**-—**) vs with baseline control parameter values (- - -) when feed parameters were optimized.

#### 3.4.1 Methanogen Analysis

The sensitivity analysis studied how small changes in model inputs affected methane production. To investigate how larger changes in parameter values affect methane production, the model was evaluated at varying initial methanogen abundances to investigate the possibility of reducing methane production by reducing methanogen abundance.

Table 5 shows methane output changes depending on initial methanogen abundance. Greater reduction in methane is seen at lower methanogen abundances, but some reductions can be seen already at 25% of control baseline methanogens. This illustrates that methane output is not very sensitive to methanogen abundance in control conditions until methanogen abundance is decimated. The sensitivity to methanogen abundance at low levels of methanogens was not seen in the sensitivity analysis, as the local and global sensitivity analysis had only varied parameters 50 -150% of baseline values.

**Table 5:** Methane production at various methanogen abundances.

| % of Control Methanogens | Final Methane Output | % Reduction in Methane |
| --- | --- | --- |
| 100 | $4.37 \cdot 10^{-2}$ | 0 |
| 75 | $4.35 \cdot 10^{-2}$ | 0.46 |
| 50 | $4.29 \cdot 10^{-2}$ | 1.82 |
| 25 | $3.97 \cdot 10^{-2}$ | 9.14 |
| 10 | $3.30 \cdot 10^{-2}$ | 24.5 |
| 5 | $2.84 \cdot 10^{-2}$ | 35.0 |
| 2.5 | $2.01 \cdot 10^{-2}$ | 54.0 |
| 1 | $8.30 \cdot 10^{-3}$ | 81.0 |
| .1 | $48.45 \cdot 10^{-4}$ | 98.1 |
| 0 | 0 | 100 |

The large reduction in methane output seen in Table 5 suggests the possibility of reducing methane by greatly reducing methanogen concentration. A 99% reduction in in control methanogens reduces final methane output by 98.1%.

## 4 Discussion

Microbial interactions in the rumen are highly complex, making disentangling how these interactions affect total rumen fermentation and subsequent methane production difficult. Mathematical models present one way to resolve this complexity. The application of quantitative methods to previously obtained rumen microbial data and *in silico* experimentation allows for the identification of key parameters predicted to affect methane production, which can then be further evaluated *in vitro* and *in vivo*. In addition, these *in silico* predictions reduce the need for extensive exploratory *in vitro* and *in vivo* experimentation, together saving time and money while simultaneously reducing the need for animal trials. Though mathematical modeling presents a possible method for the identification of future methane mitigation strategies, widespread application first requires the building, calibration, and validation of models able to predict microbial shifts accurately in the rumen and subsequent outcomes. In this work we calibrated a previously established rumen fermentation model with data collected from an *in vitro* trial testing the effects of promising feed additive, *A. taxiformis* (3,12).

Here, a mechanistic mathematical model of *in vitro* rumen fermentation that considers the effect of *A. taxiformis* was adapted to the Roque et al. (3) experimental setup, calibrated with virtual microbiome experiments, and validated with Roque et al. (3) microbial profiles and gas production data. The modified mathematical model satisfactorily fit both the CH_4_ and CO_2_ data from Roque et al. (3)’s *in vitro* RUSITEC experiment. The virtual microbiome experiments performed here studied the importance of functional classifications as they pertain to methane output in the rumen ecosystem. Additionally, the relationship between methanogen abundance and methane production was analyzed. The model predicts a significant reduction in final methane production in control conditions when methanogen abundance is sufficiently low. However, the complete eradication of a functional group in any microbial context is not practical *in vivo*. Rather this type of *in silico* experimentation allows for the exploration of biological bounds and can suggest routes for future exploration that would otherwise be unknown.

Local and global sensitivity analyses were used to study the predicted effects of the rumen microbial community on methane production. In doing so we were able to show that the effects of microbial variation on methane production are greater with the addition of *A. taxiformis*. Additionally, the notable effect that the inclusion of this red seaweed alters system functional distributions, is an interesting finding from the virtual experimentation. *In vivo* it is likely the addition of *A. taxiformis* has effects for the microbiome diversity and function beyond the reduction in methanogens, supporting that the newly calibrated and tested model from this work accurately reflects complex shifts in the rumen ecosystem that result from exogenous substrates in the diet (38–40).

Identifying the most sensitive parameters can motivate further experimental investigations. A better understanding of these parameter values can increase the reliability of model predictions. The sensitivity analysis identified a different set of most significant parameters in control and treatment conditions. In treatment conditions, the top two top parameters were bromoform and a parameter of the effects of bromoform on other processes. In our data we only had only one concentration of bromoform, limiting our ability to determine the effects of this compound across a range of inclusion values. Dynamic measurements of bromoform and hydrogen as well as more inclusion rates of bromoform would be helpful to have more certainty in parameter values and the resulting model predictions. Another parameter identified in the sensitivity analysis in treatment conditions is the total microbial concentration (TMC). To have more certainty in its value, it is necessary to measure total microbial dry matter or quantify absolute bacterial abundance when collecting sequencing data for modelling purposes. In control conditions, the most sensitive parameters included feed composition parameters and parameters related to their utilization. Feed composition parameters are directly controllable. We demonstrated that predicted final methane output without bromoform in the system could be reduced by 25.6% with optimal feed composition parameters. This reduction in methane is predicted when initial neutral detergent fiber concentration is reduced by 49.7%, initial nonstructural carbohydrate concentration is reduced by 38.0%, and initial protein concentration is increased by 49.6%.

The objective of this work was to adapt and calibrate the Muñoz-Tamayo et al. (12) mathematical model of *in vitro* rumen fermentation with the Roque et al. (3) gas and microbial phylogenetic data from the *in vitro* semi-continuous RUSITEC system and conduct sensitivity analysis and identify rumen parameters that might be key drivers in enteric methane production. Mathematical models, like Muñoz-Tamayo et al. (12), are powerful tools as they allow tracking of individual state variables during rumen fermentation simulations. The confirmation of the model’s ability to predict changes which reflect known *in vivo* phenomena such as microbial community shifts in response to feed changes suggests the ability to utilize such models for the investigation of systemic effects of other feed additives prior to *in vivo* or even *in vitro* experimentation. While this modelling approach effectively captures methane and carbon dioxide production, additional data would help enhance our understanding of the complete rumen microbiome dynamics. Dynamic measurements of bromoform and hydrogen as well as more inclusion rates of bromoform are necessary to further validate the model. Additionally, model extensions could account for other feed additives and *in vivo* conditions, making this rumen fermentation model an even more useful tool for the field at large.

## 6 Supplement

### Supplement A: Model equation details

Below we outline the Muñoz-Tamayo et al(12) model that was adapted to the Roque et al(3) experiment.

In addition to the model adaptions of the Muñoz-Tamayo et al(12) model outlined in section 2.2, our implementation of the model is slightly different. Some variables are discontinuous due to the experimental setup in Roque et al (3). In the experiment, each vessel was opened every 24 hours, a feedbag and the gas bag were exchanged, and the vessels were flushed with nitrogen before being reattached.

Thus, all gaseous phase variables (moles of hydrogen in the gas phase (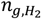 (mol)), moles of methane in the gas phase (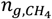 (mol)), and moles of carbon dioxide in the gas phase (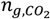 (mol))) were set to zero every 24 hours to account for the exchange of gas bags. Additionally, variables determined by feed composition of the feedbags (the neutral detergent fiber concentration (*z*_*ndf*_ (g/L)), non-structural carbohydrate concentration (*z*_*nsc*_ (g/L)), protein concentration (*z*_*pro*_ (g/L)), and bromoform concentration (*S*_*br*_ (g/L))), were adjusted every 24 hours to account for the contents of the new feedbags. To account for these discontinuities, integration was stopped at the time points of discontinuity (24 hour, 48 hour, 72 hour), initial conditions were set, and integration was restarted. Initial conditions of the gaseous phase and feed variables were set as described above and the initial conditions of the remaining variables were set to the last value of the previous integration of their variable.

### Volume and Dilution

750 milliliters rumen fluid with and without *A. taxiformis* were divided into 3 vessels each. Buffer was delivered at 0.39 mL/min to each vessel throughout days 1-4. The volume increased at 0.39 ml/min until it reached 1 liter at hour 10.68. After the vessels reach reached full capacity at 1 liter, buffer continued to be added and excess fluid overflowed. It is assumed that the fluid is well-mixed. The change in volume is described by the following ordinary differential equation:

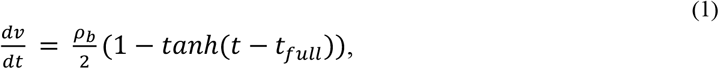

where ρ_*b*_ is the rate of saliva buffer (0.39 ml/min) and *t*_*full*_ is the time at which the vessels reach full capacity (10.68 h).

Because of the continuous addition of buffer, each state variable in solution includes a reduction in concentration because of dilution of the form:

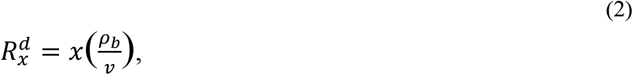

where *x* is the state variable in solution and *v* is the volume of the solution.

### Polymer Components

Polymer components in the model include neutral detergent fiber concentration (*z*_*ndf*_ (g/L)), non-structural carbohydrate concentration (*z*_*nsc*_ (g/L)), and protein concentration (*z*_*pro*_ (g/L)). Neutral detergent fiber, non-structural carbohydrates, and proteins are hydrolyzed at rates of *ρ*_*ndf*_, *ρ*_*nsc*_, and *ρ*_*pro*_ respectively. The equations for the hydrolysis rates are the product of hydrolysis rate constants and the concentration of their polymer:

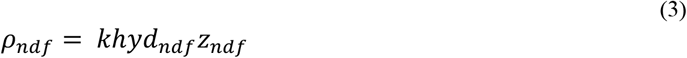

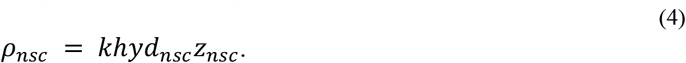

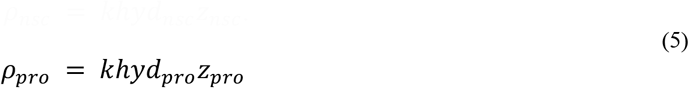

Sugar utilizers, amino acid utilizers and methanogens die at rates of 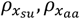, *and* 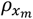which are defined in equation (21), (22), and (23). Dead microbial cells become a source of non-structural carbohydrates and proteins at the rate of the product of molecular weight of microbial cells (*w*_’A_), the rate of microbial death (*ρ*_*xsu*_ + *ρ*_*xaa*_ + *ρ*_*xm*_), and the fraction of carbohydrates (σ_=*ch,x*_) and proteins (*σ*_*pro,x*_) in the biomass, respectively. The ordinary differential equations that describe the change of the polymer component concentrations are as follows:

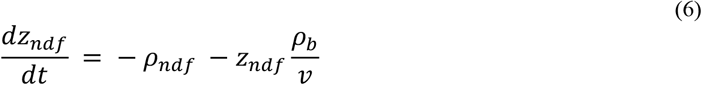

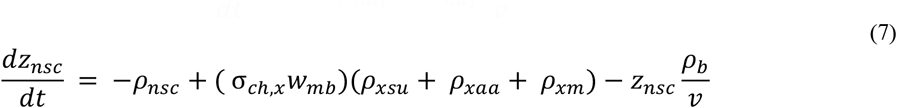

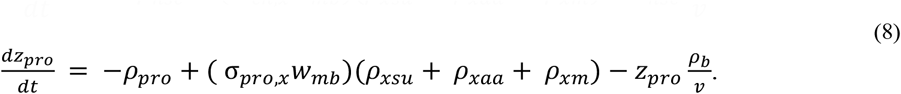

### Monomer Components

The model describes the change sugar concentration (*S*_*su*_ (mol/L)) and average amino acid concentration (*S*_*aa*_ (mol/L)).

The model assumes that sugar and amino acid monomers are released through the hydrolysis of neutral detergent fiber, non-structural carbohydrate, and protein. The hydrolysis rate of the polymers is divided by the respective molecular weight of the monomer because sugar and amino acid concentration are tracked in mol/L and neutral detergent fiber, non-structural carbohydrates, and proteins are tracked in g/L. Sugar is released through the hydrolysis of fiber and non-fiber carbohydrates at a rate of 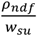 and 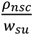 respectively, where *w* is the molecular weight of sugar.

Amino acid is released through the hydrolysis of proteins at a rate of 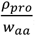, where *w*_*aa*_ is the molecular weight of amino acid.

Amino acids and sugars are utilized at a rate of *ρ*_*aa*_ and *ρ*_*su*_, respectively. *ρ*_*aa*_ is defined as

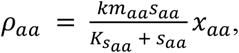

where *km*_*aa*_ is the maximum specific utilization rate constant of amino acids, 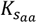 is the Monod constant associated with the utilization of amino acids, and *x*_*aa*_ is the concentration of amino acid utilizers.

*ρ*_*su*_ is defined as

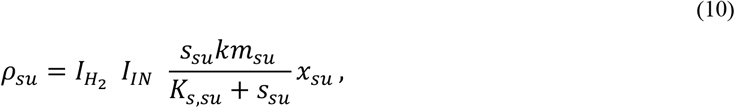

where *I*_*IN*_ is the nitrogen limitation factor is defined as

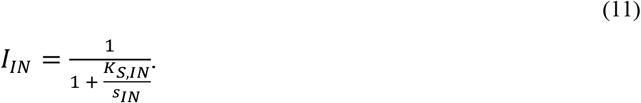

*K*_*S,IN*_ is the nitrogen limitation constant and *S*_*IN*_ is inorganic nitrogen and defined in equation (41).

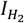 is the hydrogen control factor and is defined as

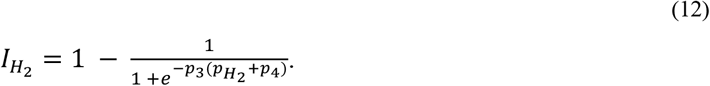

*p*_3_and *p*_4_ are both parameters which were estimated in Muñoz-Tamayo et al(12) to be 102.5300 and - 0.2350 respectively. 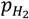 is the partial pressure of hydrogen and is defined as

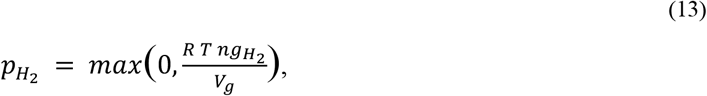

where R is the ideal gas constant, T is temperature, and *V*_*g*_ is the volume in gas phase which is defined in equation (31).

The ordinary differential equations that describe the change of the polymer component concentrations are as follows:

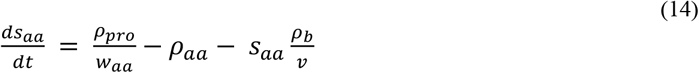

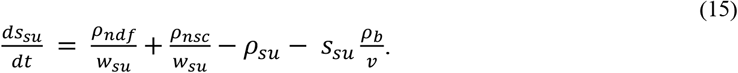

### Functional Microbial Groups

The model describes the change of sugar utilizer concentration (*x*_*su*_ (mol/L)), amino acid utilizer concentration (*x*_*aa*_ (mol/L)), and methanogen concentration (*x*_*m*_ (mol/L)).

The functional microbial group of sugar utilizers utilizes sugar and NH_3_, the amino acid utilizers metabolize amino acids, and the methanogens convert H_2_ to produce CH_4_.

The utilization rate of hydrogen 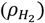 is described as follows:

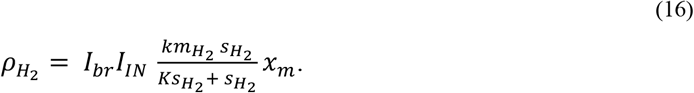

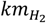 is the maximum specific utilization rate constant of hydrogen. 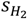 is the concentration of hydrogen in the liquid phase. 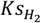 is the Monod constant associated with the utilization of hydrogen. *I*_*IN*_ is the nitrogen limitation factor and was defined in equation (11). *I*_*br*_ is the inhibition factor of the methanogens growth rate by the action of bromoform and is equal to

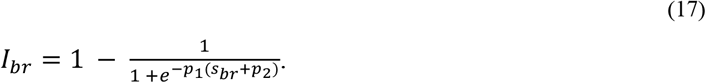

*S*_*br*_ is the concentration of bromoform and defined in equation (43). *p*_1_ and *p*_2_ effect the inhibition factor of methanogens growth rate by the action of bromoform and were estimated.

The utilization rate of amino acids and sugars are *ρ*_*aa*_ and *ρ*_*su*_ and are defined in equation (9) and (10)respectively.

The growth rates of sugar utilizers, amino acid utilizers, and methanogens are:

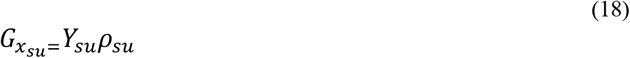

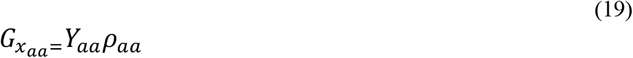

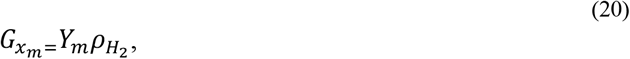

where *Y*_*i*_ is the microbial biomass yield factor of functional group i.

The cell death rates are:

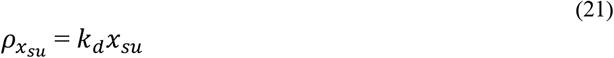

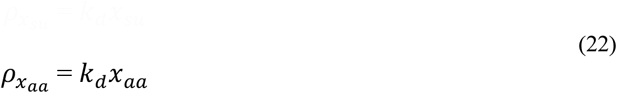

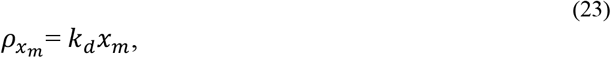

where *k*_*d*_ is the death cell rate constant for all microbial groups.

The ordinary differential equations that describe the change of the functional microbial group concentrations are as follows:

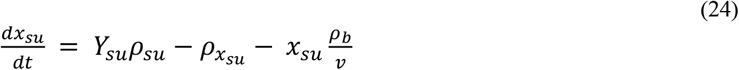

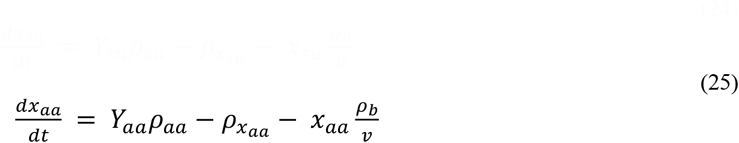

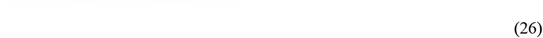

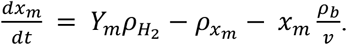

### Volatile Fatty Acids

Volatile fatty acids in the model include acetate concentration (*S*_*ac*_ (mol/L)), butyrate concentration (*S*_*bu*_ (mol/L)), and propionate concentration (*S*_*pr*_ (mol/L)). In addition to microbial growth, the utilization of sugar and amino acids produce products, such as volatile fatty acids. *Y*_*i, j*_ is the yield factor of component i during utilization of substrate j and is explained in detail in the yield factor calculation section.

The ordinary differential equations that describe the change of the volatile fatty acid concentrations are as follows:

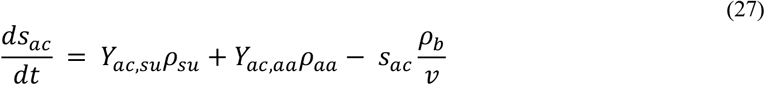

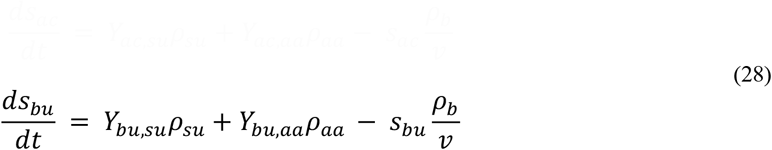

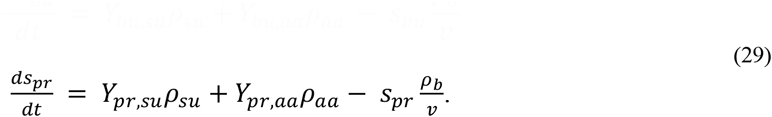

#### Hydrogen, Methane, and Carbon Dioxide

During the fermentation process, hydrogen, methane, and carbon dioxide are produced. Liquid-gas transfer occurs for all three molecules. The model tracks moles of hydrogen in the gas phase (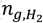 (mol)), hydrogen concentration in the liquid phase (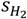 (mol/L), moles of methane in the gas phase (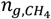 (mol)), methane concentration in the liquid phase (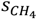 (mol/L)), and moles of carbon dioxide in the gas phase (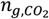 (mol)). The concentration of carbon dioxide in the liquid phase 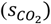 is not a state variable in the model (defined in equation 37) as it is accounted for in the state variable for inorganic carbon concentration (see equation 42)

The liquid-gas transfer rate of component i is of the form

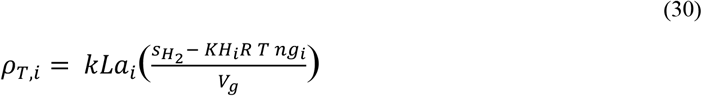

where *KH*_*i*_ is Henry’s law coefficient of component i, *R* is the ideal gas constant, *T* is the temperature, *V*_*g*_ is the volume in gas phase and defined as:

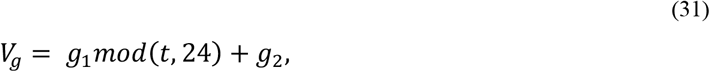

where *g*_1_ and *g*_2_ were calculated calculated using MATLAB’s function polyfit for volume data points vs time. *g*_1_ is 0.015924 and 0.0079877 and *g*_2_ is 0.054785 and 0.035973 for control and treatment respectively. Note that *g*_2_ is not zero since the system is flushed with nitrogen (i.e. there is never zero gas)

The liquid-gas transfer rates of hydrogen 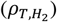 methane 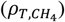, and carbon dioxide 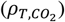 are defined as follows:

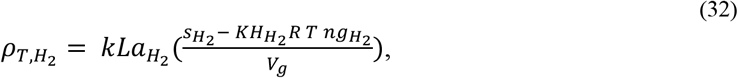

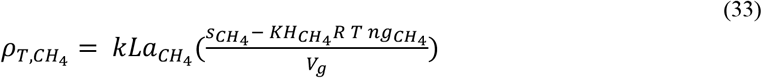

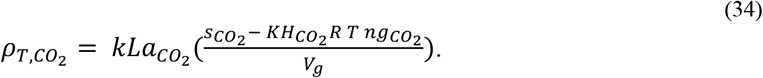

Muñoz-Tamayo et al(12) used the same liquid-gas transfer constant for *H*_2_, *CO*_2_, and *CH*_4_. We added a separate liquid-gas transfer constant for CO_2_ and H_2_.

Hydrogen in the liquid phase is produced in both sugar and amino acid utilization and utilized by methanogens. Methane in the liquid phase is produced by methanogens which utilize hydrogen.

*Y*_*i,j*_ is the yield factor of component i during utilization of substrate j and is explained in detail in the yield factor calculation section.

The ordinary differential equations that describe the change of hydrogen and methane concentration are as follows:

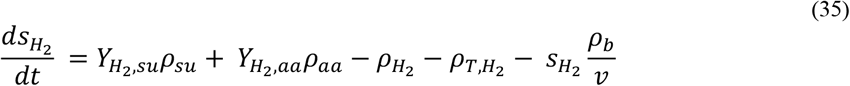

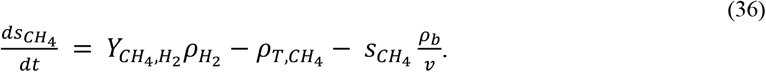

The concentration of carbon dioxide in the liquid phase is defined as

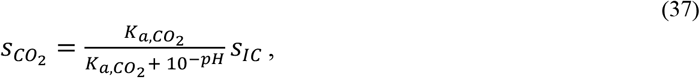

where 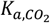 is the equilibrium constant of carbon dioxide and *S*_*IC*_ is inorganic carbon concentration (described in detail in inorganic nitrogen and inorganic carbon section).

The ordinary differential equations that describe the change of the components in the gas phase are as follows:

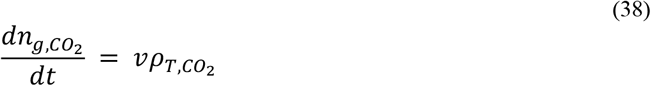

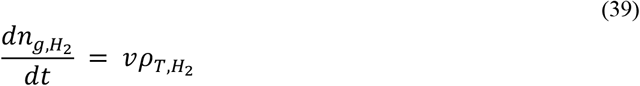

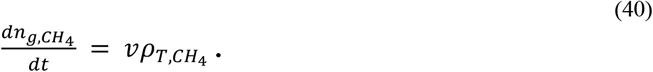

### Inorganic Nitrogen and Inorganic Carbon

Nitrogen and carbon are both necessary for microbial biosynthesis. The model tracks inorganic nitrogen concentration (*S*_*IN*_ (mol/L) and inorganic carbon concentration (*S*_*IC*_ (mol/L)). *Y*_*i,j*_ is the yield factor of component i during utilization of substrate j and is explained in detail in the yield factor calculation section.

Inorganic nitrogen is the sum of ammonia and ammonium ion concentrations. Inorganic carbon is the sum of soluble carbon dioxide and bicarbonate ion concentrations. Instead of modelling both, of soluble carbon dioxide and bicarbonate ion concentrations are gathered in the inorganic carbon variable, and ammonia and ammonium ion concentrations are gathered in the inorganic nitrogen variable.

The ordinary differential equations that describe the change of the inorganic nitrogen concentration and inorganic carbon concentration are as follows:

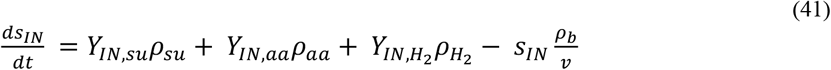

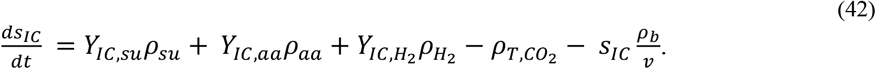

### Bromoform

#### Asparagopsis taxiformis

which contains bromoform, was added to each treatment feedbag at a 5% inclusion rate. The model assumes that bromoform concentration decays at the rate *k*_*br*_, the kinetic rate constant of bromoform utilization.

The ordinary differential equation that describes the change of the bromoform concentration is the following:

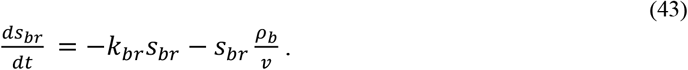

The model assumes that bromoform affects some of the model interactions through direct inhibition of methane production and indirectly through hydrogen control. The following details explain the influences of bromoform on these processes.

The rate of liquid hydrogen utilization is affected by *I*_*br*_ (inhibition factor of the methanogens growth rate by the action of bromoform). The inhibition is represented as a function of bromoform concentration. As bromoform increases, *I*_*br*_ decreases, which decreases the rate of hydrogen utilization 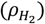. Liquid hydrogen is utilized by methanogens (*x*_*m*_) and yields methane in the liquid phase 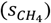, nitrogen (*S*_*IN*_), and inorganic carbon (*S*_*IC*_).

Since less *H*_2_ is utilized, there is a built-up of *H*_2_. The built-up of *H*_2_ in the liquid phase increases moles of hydrogen in the gas phase 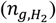. The increase of hydrogen in the gas phase 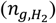, increases the partial pressure of 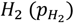, which increases the hydrogen control factor for sugar utilization 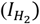. An increase in the hydrogen control factor for sugar utilization increases the utilization rate of sugars (*ρ*_*su*_), which increases the concentration of sugar utilizing microbes, total acetate concentration (*S*_*ac*_), total butyrate concentration (*S*_*bu*_), and total propionate concentration (*S*_*pr*_).

### Yield Factor Calculation

Next, we describe how yield factors were calculated. Muñoz-Tamayo et al(12) used stoichiometry of fermentation to determine yield factors using well known reactions. They assumed that the molecular formula for microbial biomass is C_5_H_7_O_2_N. The reactions considered are outlined below.

#### Sugar Utilization

Reaction 1: C_6_H_12_O_6_ + 2H_2_O → 2CH_3_COOH +2CO_2_ + 4H_2_

Reaction 2: 3C_6_H_12_O_6_ → 2CH_3_COOH + 4CH_3_CH_2_COOH + 2CO_2_ +2H_2_O

Reaction 3: C_6_H_12_O_6_ → CH_3_CH_2_CH_2_COOH + 2CO_2_ + 2H_2_

Reaction 4: 5C_6_H_12_O_6_ + 6NH_3_ → 6C_5_H_7_O_2_N + 18H_2_O

Reaction 1 to Reaction 4 describe the utilization of sugars.

λ_1_, λ_2_, and λ_3_ are the fract*i*on of *su*gar *u*t*i*l*i*zed v*i*a react*i*on 1, 2, or 3 respect*i*vely.

λ_1_, λ_2_ and λ_3_ effect various yield factors, including the yield factor of acetate (*ac*), propionate (*pr*), butyrate (*bu*), hydrogen (*H*_2_), inorganic carbon (*IC*), and inorganic nitrogen (*IN*) during sugar utilization. λ_1_, λ_2_, and λ_3_ are defined as follows:

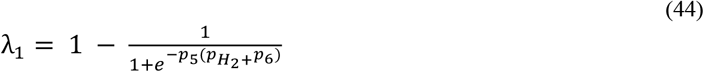

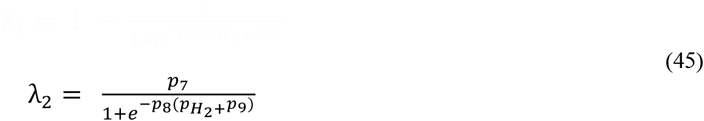

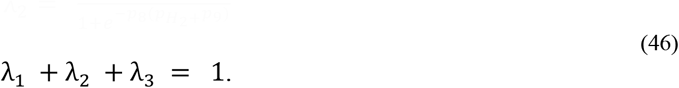

The four parameters *p*_5_, *p*_6_, *p*_7_, *p*_8_, and *p*_9_ were estimated in Muñoz-Tamayo et al(12) as 7.5020, 0.0698, 0.9219, 2.5533, and 0.1675 respectively.

The fraction of glucose utilized in catabolism (*fSu*) based on react*i*on 4 *i*s

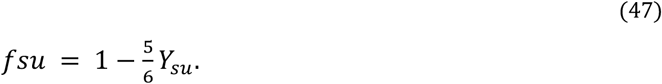

The constant *Y*_*su*_ is the microbial biomass yield factor of sugar utilizers.

*Y*_*i,j*_ is the yield factor of component i during utilization of substrate j. The yield factors from sugar utilization are as follows:

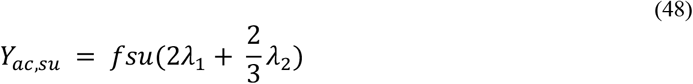

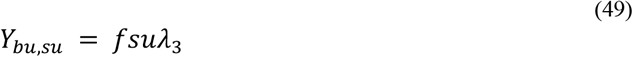

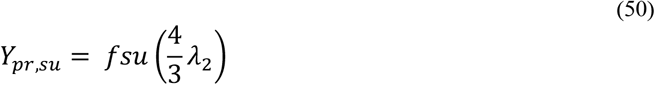

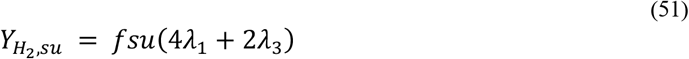

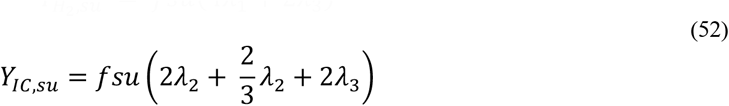

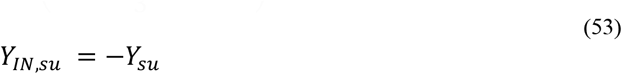

#### Amino Acid Utilization

Reaction 5: C_5_H_9_._8_O_2·7_N_1.5_ → (1™Y_aa_)·*σ*_ac,aa_CH_3_COOH

+ (1™Y_aa_) ·*σ*_pr,aa_CH_3_CH_2_COOH + Y_IN,aa_NH_3_

+ (1™Y_aa_) · *σ*_bu,aa_CH_3_CH_2_CH_2_COOH+(1™Y_aa_) ·*σ*_IC,aa_CO_2_

+(1™Y_aa_) · *σ*H_2,aa_H_2_ +Y_aa_ C_5_H_7_O_2_N

Reaction 5 describes the utilization of amino acids. *Y*_*aa*_ is the microbial biomass yield factor of amino acid utilizers. The yield factors from amino acid utilization are as follows:

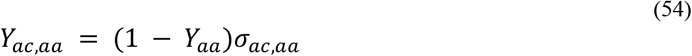

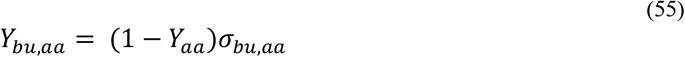

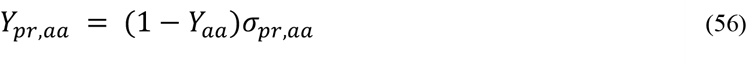

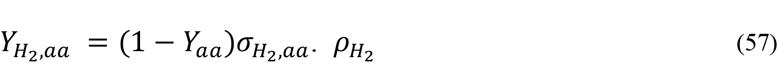

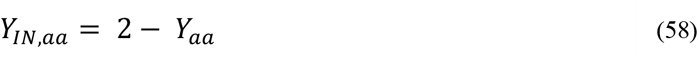

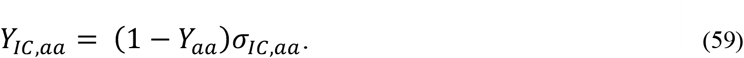

### Hydrogen Utilization

Reaction 6: 4H_2_ +CO_2_ →CH_4_ +2H_2_O

Reaction 7: 10H_2_ + 5CO_2_ + NH_3_ → C_5_H_7_O_2_N + 8H_2_O

Reaction 6 and 7 describe hydrogen utilization. *fH*_2_ is the fraction of *H*_2_ utilized in catabolism and is defined as follows:

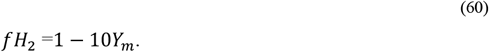

*Y*_*m*_ is the microbial biomass yield factor of methanogens. The yield factors from hydrogen utilization are as follows:

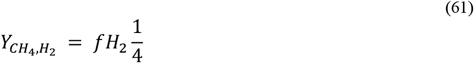

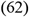

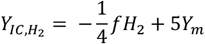

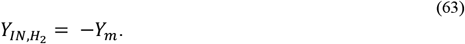

### Summary of Equations

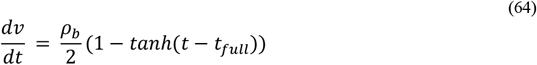

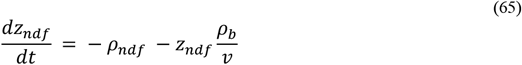

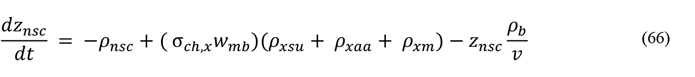

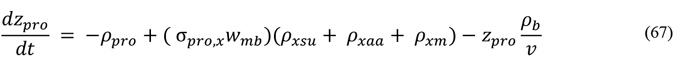

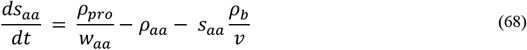

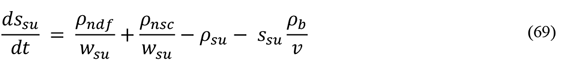

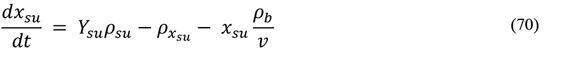

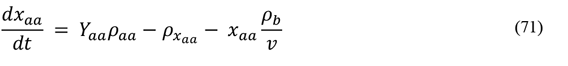

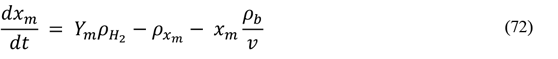

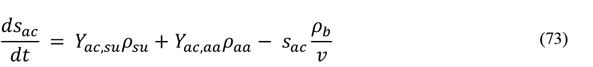

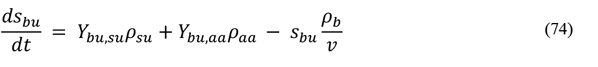

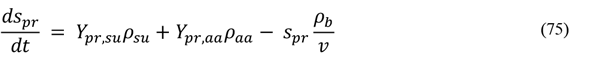

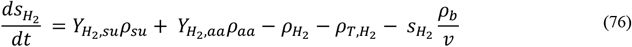

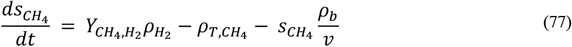

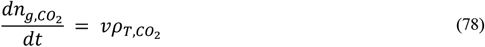

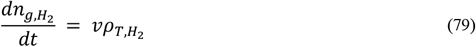

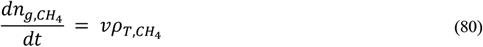

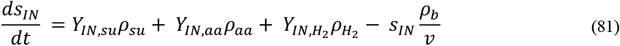

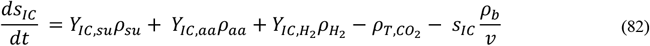

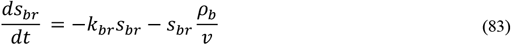

#### Supplement B: Secondary Parameter Estimation and Sensitivity Analysis

A secondary parameter estimation was conducted with the microbial functional concentration initial conditions from the virtual microbiome experiments. It shows that model fit does not significantly improve with an additional parameter estimation. Additionally, a sensitivity analysis with this parameter set identified a similar set of most significant parameters.

Model parameters were estimated with MATLAB’s optimization toolbox constrained nonlinear optimization solver **fmincon** with default values for termination tolerances. Parameters were varied over the same relative bounds as the initial parameter estimation to minimize the cost function of the sum of the squared differences between model CH_4_ and CO_2_ output and CH_4_ and CO_2_ data for 96 hours for control and treatment conditions.

Parameters that were estimated, nominal values, and the relative difference can be found in S1 Table. Estimated parameters include 17 parameters varied separately for control and treatment conditions, two parameters estimated only in treatment conditions (the kinetic rate constant of bromoform utilization, and the initial bromoform concentration), and one parameter estimated together for control and treatment conditions (total microbial concentration).

S1 Figure shows the model fits of CH_4_ and CO_2_ concentrations from RUSITEC samples generated with and without *A. taxiformis* added to the feed. The difference of parameter values from the initial compared to the secondary parameter estimation can be found in S1 Table. The average relative difference was 21.2%. Only one parameter, the hydrolysis rate constant of cell wall carbohydrates, had a relative difference greater than 50% and was identified as significant in the sensitivity analysis.

To quantitatively assess the accuracy of the fits, we performed a linear fit of prediction as a function of observation, to describe the relationship between the data and model output. The linear fitted curve was calculated using the least squares method, by minimizing the sum of the square of errors between the data and the model output. The smaller the absolute value of the y-intercept and the closer the slope is to one, the better the model prediction. The observed data plotted against the prediction of the fits, as well as the calculated best fit line CH_4_ and CO_2_ are shown in S2 Figure. The linear regression equations of control and treatment CH_4_ and control and treatment CO_2_ had slopes of 1.00, 0.83,1.08, and 1.27 respectively and all had y-intercepts with absolute value than .004. This shows that the model satisfactorily captures the dynamical behavior of both CH_4_ and CO_2_ after the secondary parameter estimation and is comparable to the fits after the initial parameter estimation and after the virtual microbiome experiments.

A sensitivity analysis was then performed with this new parameter set (parameters included are indicated with a * in Table 1).

We examined the sensitivity of the model’s output of the final gaseous methane for two conditions:

1. Control: Roque et al. (3) control experimental conditions and parameter values that were estimated for the control conditions during the second parameter estimation.
2. Treatment: Roque et al. (3) treatment experimental conditions and parameter values that were estimated for treatment conditions during the second parameter estimation.

The coefficient of variation in the final methane output is 0.26 and 1.11 for control and treatment respectively. The coefficient of variation is the standard deviation divided by the mean and is provided as a scale for the Sobol indices between conditions. It is significantly higher for treatment, similarly to the initial sensitivity analysis.

The results of the global sensitivity analysis for each parameter whose total order Sobol index was greater than 0.05 for control or treatment conditions for control and treatment can be found in S3 Figure.

The total order Sobol index is greater than 0.05 for seven control and five treatment parameters. The selected parameter sets include one common parameter, the maximum specific utilization rate of hydrogen (km_h2_). Additionally, control included the three feed initial conditions (z_ndf0_, z_nsc0_, z_pro0_), one hydrolysis rate constant (khyd_ndf_), a parameter affecting the maximum of a sigmoid function of the fraction of sugar utilization via reaction 2 (p_7_; see Supplement A for details), and the microbial biomass yield factor of sugar utilization (Y_su_). Treatment included the total microbial concentration (TMC), the initial percent abundance of methanogens (p_m0_), the maximum specific utilization rate of hydrogen (km_h2_), a parameter affecting the steepness of a sigmoid function of the inhibition factor of the methanogens growth rate by the action of bromoform (p_1_; see Supplement A for details), and initial bromoform concentration (s_br0_). The only additionally identified parameters in the secondary sensitivity analysis are the microbial biomass yield factor of sugar utilization (Y_su_) and the maximum specific utilization rate of hydrogen (km_h2_) in the control condition. The parameters contributing most to overall variance did not change in the secondary sensitivity analysis.

**S1 Figure:**
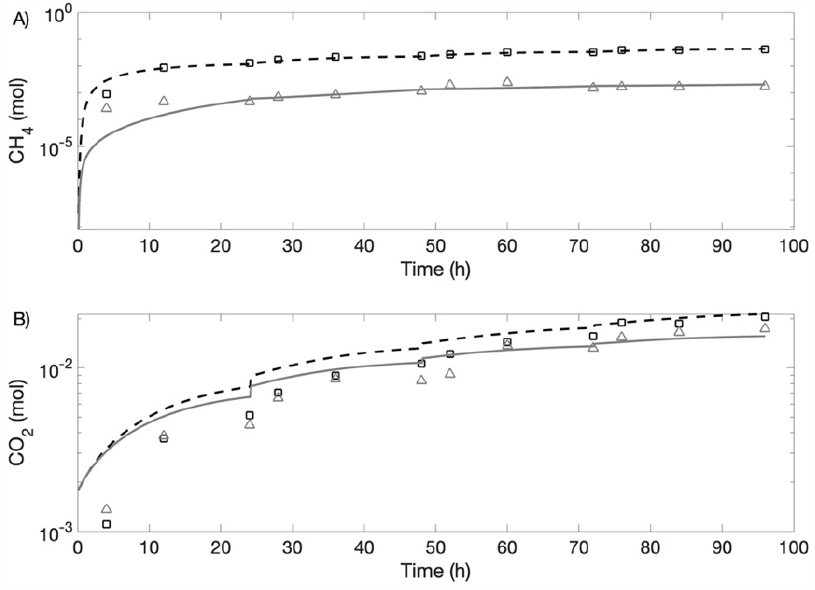
Model calibration with experimental data after secondary parameter estimation. Experimental data of treatment (square) and control (triangle) are compared against model response of treatment (**- - -**) and control (**-—**) for CH_4_ and CO_2_. (A) Control. (B) Treatment

**S2 Figure:**
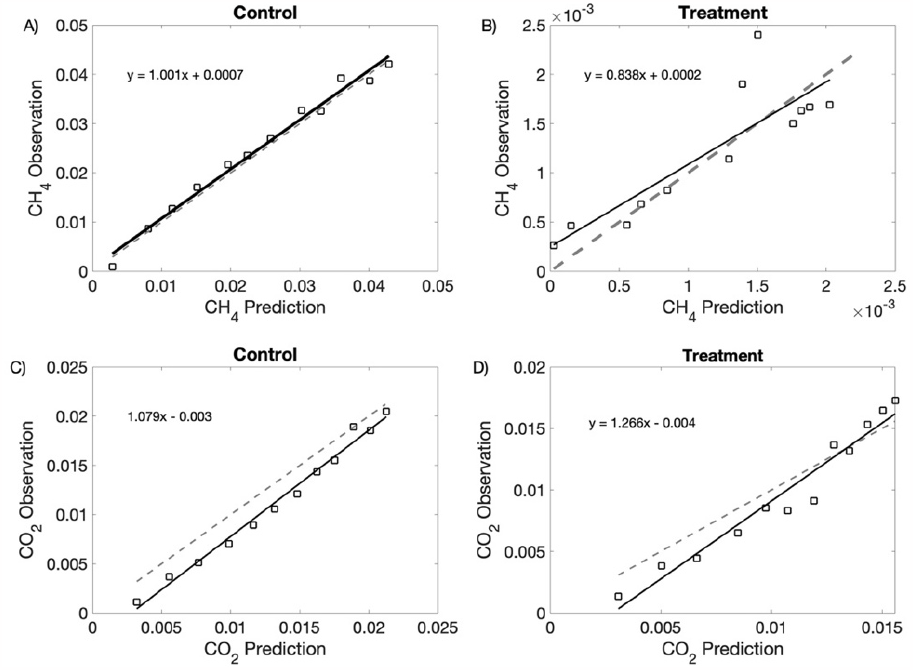
Observations plotted against model predictions (square) after secondary parameter estimation. Linear fitted curve is the solid black line (equation of line displayed on graph). Line y = x is the gray dashed line. The linearity shows the reliability of the model.

**S3 Figure:**
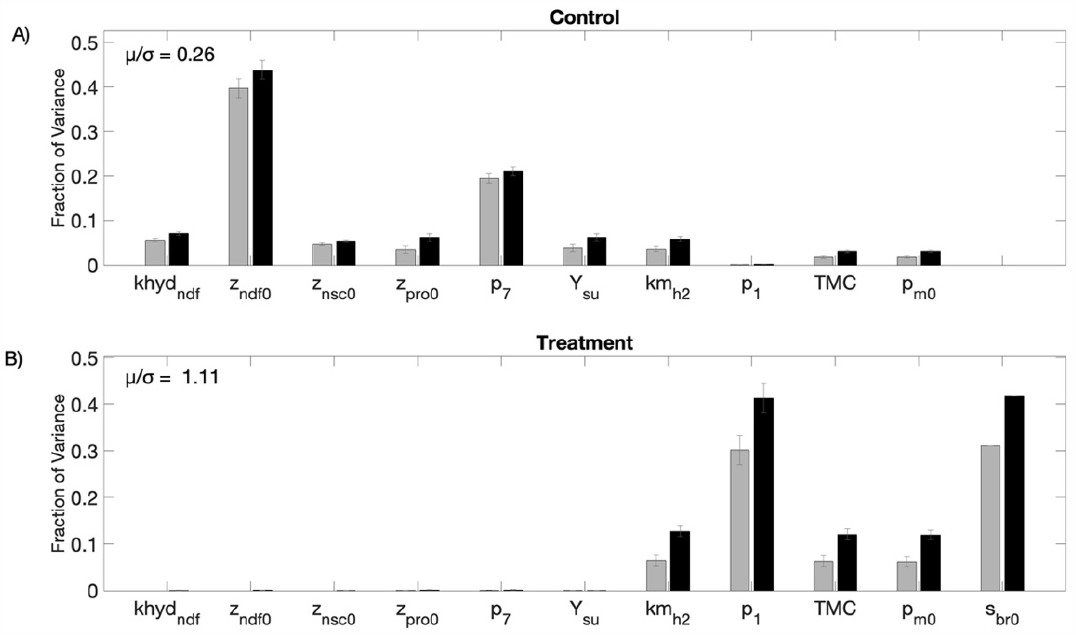
Sensitivity Analysis after secondary parameter estimation. *Global Sensitivity Analysis:* First (gray) and Total (black) Order Sobol indices are plotted as bars with errors of 95% confidence interval. The coefficient of variation (μ/σ) is displayed, for comparing the fraction of variance between conditions.

**S1 Table:** Estimated values after secondary parameter estimation.

| Parameter | Estimated Value |  | Nominal Value |  | Relative Difference |  |
| --- | --- | --- | --- | --- | --- | --- |
|  | Control | Treatment | Control | Treatment | Control | Treatment |
| initial $z_{ndf}$ | 6.45 | 5.18 | 6.13 | 5.13 | 5.3% | 1.0% |
| initial $z_{nsc}$ | 2.04 | 1.83 | 2.02 | 1.86 | 13.8% | 1.4% |
| initial $z_{pro}$ | 2.82 | 3.06 | 3.14 | 2.97 | 5.4% | 3.2% |
| initial $s_{br,0}$ | NA | $3.07 \cdot 10^{-3}$ | 0 | $3.74 \cdot 10^{-3}$ | N/A | 17.9% |
| $TMC$ | $1.87 \times 10^{-2}$ | $1.87 \times 10^{-2}$ | $3.37 \times 10^{-2}$ | $3.37 \times 10^{-2}$ | 44.3% | 44.3% |
| $k_{br}$ | N/A | $1.15 \cdot 10^{-3}$ | N/A | $1.68 \cdot 10^{-3}$ | N/A | 31.7% |
| $k_{hyd,ndf}$ | $7.76 \cdot 10^{-2}$ | $2.90 \cdot 10^{-2}$ | $4.80 \cdot 10^{-2}$ | $4.09 \cdot 10^{-2}$ | 61.6% | 29.0% |
| $k_{hyd,nsc}$ | $1.08 \cdot 10^{-1}$ | $1.09 \cdot 10^{-1}$ | $1.46 \cdot 10^{-1}$ | $8.08 \cdot 10^{-2}$ | 26.0% | 34.7% |
| $k_{hyd,pro}$ | $3.78 \cdot 10^{-1}$ | $1.05 \cdot 10^{-1}$ | $1.66 \cdot 10^{-1}$ | $1.31 \cdot 10^{-1}$ | 77.1% | 20.0% |
| $k_{m,aa}$ | 2.20 | 2.23 | 2.58 | 2.08 | 15.0% | 7.3% |
| $k_{m,H_2}$ | 16.18 | 15.26 | 16.26 | 15.89 | 0.5% | 20.6% |
| $k_{m,su}$ | 2.24 | $9.42 \cdot 10^{-1}$ | 2.00 | $9.32 \cdot 10^{-1}$ | 12.2% | 1.0% |
| $Y_{aa}$ | $2.49 \cdot 10^{-2}$ | $5.74 \cdot 10^{-2}$ | $1.61 \cdot 10^{-2}$ | $6.43 \cdot 10^{-2}$ | 54.9% | 10.7% |
| $Y_m$ | $2.00 \cdot 10^{-3}$ | $4.43 \cdot 10^{-4}$ | $3.87 \cdot 10^{-3}$ | $1.21 \cdot 10^{-3}$ | 48.2% | 63.5% |
| $Y_{su}$ | $1.66 \cdot 10^{-1}$ | $9.90 \cdot 10^{-2}$ | $1.04 \cdot 10^{-1}$ | $2.25 \cdot 10^{-2}$ | 59.7% | 56.0% |
| $kLa_{CH_4}$ | 4.41 | 8.40 | 4.58 | 8.41 | 37.4% | 0.1% |
| $kLa_{CO_2}$ | 4.06 | 7.79 | 4.07 | 8.25 | 0.3% | 5.5% |
| $kLa_{H_2}$ | 4.25 | 7.98 | 4.40 | 8.25 | 34.3% | 3.3% |
| $p_1$ | 1694.6 | 1694.6 | 1694.6 | 1694.6 | 0% | 0% |
| $p_2$ | $3.51 \cdot 10^{-4}$ | $2.603 \times 10^{-5}$ | $2.14 \cdot 10^{-4}$ | $2.21 \times 10^{-5}$ | 64.2% | 8.0% |

## Acknowledgments

This work was supported in part by NSF DMS Postdoctoral Fellowship award 2103380 to KGL.

